# The unfolded protein response triggers the immune deficiency pathway in ticks

**DOI:** 10.1101/2021.08.31.458430

**Authors:** Lindsay C. Sidak-Loftis, Kristin L. Rosche, Natasha Pence, Jessica K. Ujczo, Joanna Hurtado, Elis A. Fisk, Alan G. Goodman, Susan M. Noh, John W. Peters, Dana K. Shaw

## Abstract

The insect immune deficiency (IMD) pathway is a defense mechanism that senses and responds to Gram negative bacteria. Ticks lack genes encoding upstream components that initiate the IMD pathway. Despite this deficiency, core signaling molecules are present and functionally restrict tick-borne pathogens. The molecular events preceding activation remain undefined. Here, we show that the Unfolded Protein Response (UPR) initiates the IMD network in *Ixodes scapularis* ticks. The endoplasmic reticulum (ER) stress receptor, IRE1α, is phosphorylated in response to tick-borne bacteria, but does not splice the mRNA encoding XBP1. Instead, through protein modeling and reciprocal pulldowns, we show that *Ixodes* IRE1α complexes with TRAF2. Disrupting IRE1α-TRAF2 signaling blocks IMD pathway activation and diminishes the production of reactive oxygen species. Through *in vitro*, *in vivo,* and *ex vivo* techniques we demonstrate that the UPR-IMD pathway circuitry limits the Lyme disease-causing spirochete *Borrelia burgdorferi* and the rickettsial agents *Anaplasma phagocytophilum* and *A. marginale* (anaplasmosis). Altogether, our study uncovers a novel linkage between the UPR and the IMD pathway in ticks.

## INTRODUCTION

Arthropod-borne diseases continue to be a substantial source of morbidity and mortality worldwide^1^. Factors influencing the ability of arthropods to harbor and transmit pathogens are incompletely understood, although progress on this front has been made in recent years. Arthropod immunity is an important force in shaping vector competency^2–8^. For example, humoral defense networks such as the Immune Deficiency (IMD) pathway recognize and restrict invading microbes. As classically defined in *Drosophila melanogaster*, IMD pathway signaling events are similar to the tumor necrosis factor receptor (TNFR) pathway in mammals, but instead respond to the Gram negative bacterial PAMP (pathogen-associated molecular patterns), DAP (diaminopimelic acid)-type peptidoglycan^9, 10^. Pathway initiating receptors PGRP-LC and PGRP-LE (peptidoglycan recognition proteins LC and LE) recruit adapter molecules IMD and FADD (fas-associated protein with death domain)^11, 12^, the latter pairing with DREDD (death-related ced-3/Nedd2-like protein)^13^ which cleaves IMD. The E3 ubiquitin ligase IAP2 (inhibitor of apoptosis 2) and E2 conjugating enzymes Bendless, Uev1a, and Effette then promote (K)63 polyubiquitylation of IMD^9, 10, 14^. The resulting signaling scaffold leads to cleavage of the NF-κB signaling molecule Relish, which translocates to the nucleus and promotes antimicrobial peptide (AMP) expression^10, 14^.

Significant advances in characterizing arthropod immunity have been possible owing to the insect model organism, *Drosophila*. However, deviations from classically defined fly immunity have been reported. For example, some IMD pathway components are not found in the genomes of arachnids (ex. mites, spiders, etc.) or several hemimetabolous insects such as lice, bed bugs, psyllids, squash bugs, and whiteflies^15–27^. Triatomine bugs recently had many IMD pathway components identified, but are missing the gene encoding IMD itself^32–34^. *Ixodes scapularis* ticks lack genes encoding upstream regulators of the IMD pathway including transmembrane *PGRPs*, *imd,* and *fadd*^15, 27, 28, 30^. Despite the absence of upstream regulators, core IMD signaling molecules are active against infection^28, 30, 33–35^. Activity of the *Ixodes* IMD pathway hinges on Bendless, Uev1a, XIAP (X-linked inhibitor of apoptosis), p47, Relish, and the negative regulator Caspar, which functionally restricts tick-borne pathogens *Borrelia burgdorferi* (Lyme disease) and *Anaplasma phagocytophilum* (granulocytic anaplasmosis)^5, 28, 30, 31^. In the absence of classically defined pathway initiators, functionality of the core IMD cascade suggests that an alternative mode of activation exists.

Cellular stress responses are well-conserved across eukaryotes and respond to adverse environmental conditions, such as infection^36–45^. Herein, we demonstrate that a stress-response network, the Unfolded Protein Response (UPR), initiates the IMD pathway in *I. scapularis* ticks. *B. burgdorferi* and *A. phagocytophilum* activate the endoplasmic reticulum (ER) stress receptor IRE1α (inositol-requiring enzyme 1α), which pairs with a TRAF2-like (TNF receptor associated factor 2-like) signaling molecule (hereafter referred to as *Ixodes* TRAF2). Through molecular modeling, biochemical interactions, pharmacological manipulations, and RNAi, we show that the *Ixodes* IRE1α-TRAF2 axis functionally restricts *B. burgdorferi* and *A. phagocytophilum* in ticks, induces the IMD pathway NF-κB factor Relish, and initiates production of antimicrobial effectors. IRE1α-TRAF2 signaling also restricts the cattle pathogen *Anaplasma marginale* in *Dermacentor andersoni* ticks. Collectively, we show a fundamentally distinct mode of IMD pathway activation that explains how core signaling is activated independent of canonical upstream regulators.

## RESULTS

### The Ixodes UPR responds to tick-borne pathogens and restricts bacterial colonization

The absence of IMD pathway initiating molecules led us to hypothesize that the core signaling components may be induced through crosstalk with other molecular circuits. The UPR is a response network that is activated by ER stress through the transmembrane receptors IRE1α, PERK (PKR-like ER kinase), and ATF6 (Activating transcription factor 6). In a non-stressed state, the sensor molecule BiP (binding immunoglobulin protein) keeps all receptors inactive by binding to them^36–38^ (Fig 1A). ER stress causes BiP to disassociate from UPR receptors, allowing downstream signaling to ensue^36, 46–48^. This also results in upregulated expression of many UPR components, including BiP, with the goal of restoring cellular homeostasis ^36–38, 42, 49–51^. To evaluate whether tick-borne pathogens induce the UPR in *I. scapularis,* we quantified gene expression in *A. phagocytophilum*-infected nymphs. Relative to uninfected ticks (dotted baseline), significant increases were observed with *BiP, ire1α,* and *traf2,* suggesting that the tick UPR responds to infection (Fig 1B).

**Figure 1.**
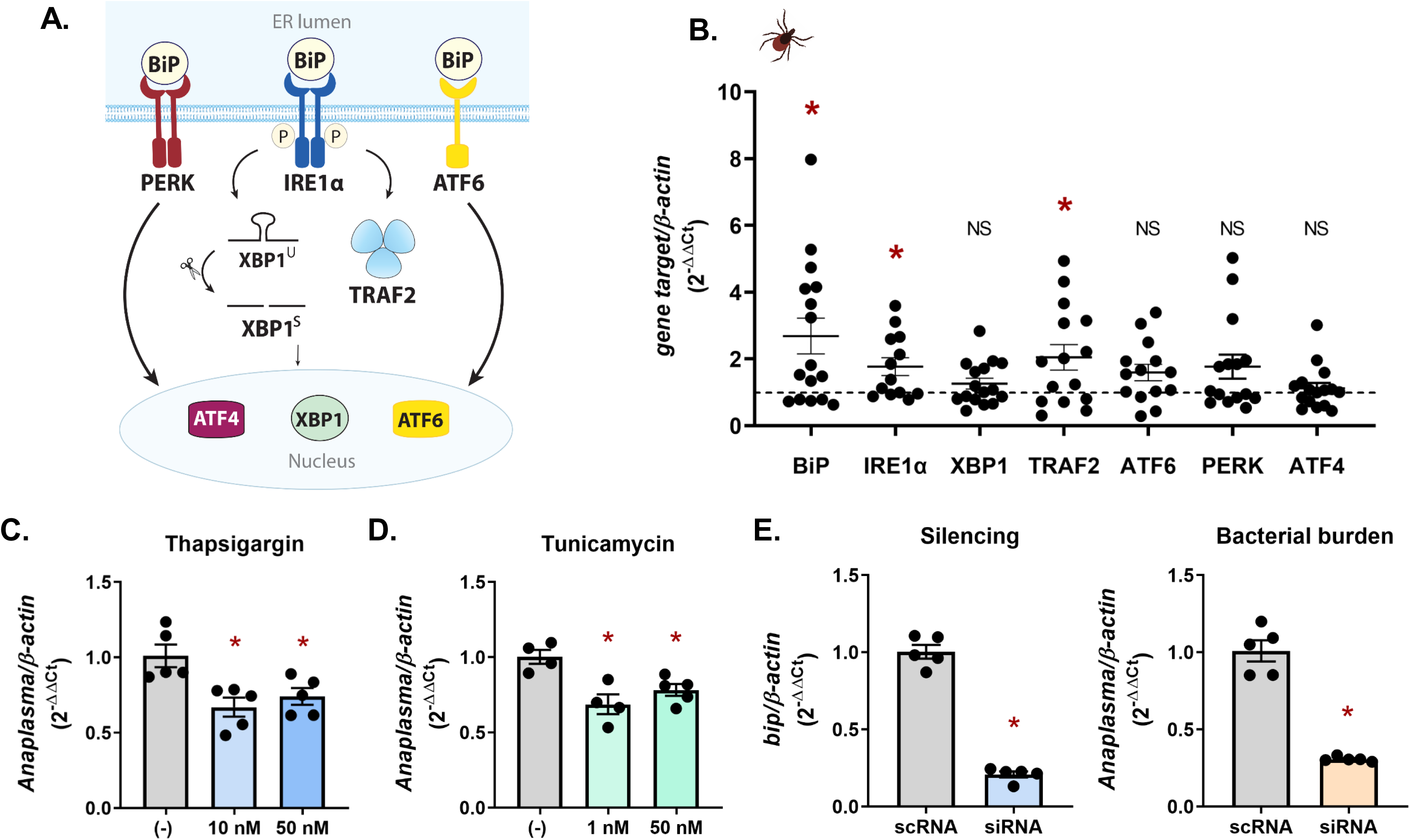
The tick UPR responds to and restricts bacterial colonization. (**a**) Graphic representation of the unfolded protein response (UPR) in mammals. (**b**) UPR gene expression in *A. phagocytophilum-*infected *I. scapularis* nymphs relative to uninfected controls (dotted line). Each point is representative of 1 nymph. Gene expression was quantified by qRT-PCR. (**c-e**) ISE6 cells (1×10^6^) were infected with *A. phagocytophilum* MOI 50 for 18 hours following a 24 hour treatment with either (**c**) thapsigargin, (**d**) tunicamycin, or (**e**) siRNA targeting the negative regulator *bip*. Gene silencing and *A. phagocytophilum* load (16s rDNA) was measured by qRT-PCR. All data shown are representative of 5 biological replicates with least two technical replicates ± SEM. Student’s t-test. *P < 0.05. scRNA, scrambled RNA; siRNA, small interfering RNA; NS, not significant. See also Supplemental Figure 1 and Supplemental Table 1.

To determine how the UPR impacts pathogen survival in ticks, we used pharmacological inducers or RNAi with the ISE6 *I. scapularis* cell line. Tick cells were treated with low doses of either thapsigargin or tunicamycin to induce ER stress prior to *A. phagocytophilum* infection. Thapsigargin inhibits the sarco/endoplasmic reticulum Ca^2+^ ATPase (SERCA), which decreases calcium levels in the ER^52^. Tunicamycin blocks N-linked glycosylation, leading to an increase of misfolded proteins^53^. Both treatments resulted in significantly less *A. phagocytophilum* (Fig 1C-D). We also used an RNAi-based approach to over activate the UPR by decreasing expression of the negative regulator BiP. In agreement with pharmacological induction, transcriptional silencing of BiP caused a decrease in *A. phagocytophilum* colonization (Fig 1E). Altogether, this demonstrates that *A. phagocytophilum* induces the UPR in ticks, which functionally restricts bacterial colonization and survival.

### Infection induces IRE1α activation, but not XBP1

Transcripts induced by *A. phagocytophilum* are associated with the IRE1α signaling axis (Fig 1A-B), which is the most conserved branch of the UPR among eukaryotes^54^. When activated, IRE1α autophosphorylates and either splices the mRNA *xbp1* (X-box binding protein 1) or signals through TRAF2^36, 37, 48, 55^ (Fig 1A). Unspliced *xbp1* mRNA (*xbp1^U^*) is held in an inactive state in the cytoplasm by forming a hairpin structure that inhibits translation. The RNase domain of IRE1α splices an internal intron from *xpb1^U^* allowing it to be translated into a protein that functions as a transcription factor^48, 56–59^ (Fig 1A). Alternatively, IRE1α can recruit the signaling molecule TRAF2 to produce proinflammatory responses through NF-κB signaling^36–38, 55^. We aligned mammalian sequences from the IRE1α pathway with tick homologs and observed sequence similarity with BiP, IRE1α, XBP1, and TRAF2 (Supplemental Figure 1A-D). Notably, the IRE1α kinase domain, RNase domain, and the activity-inducing phospho-serine (Supplemental Figure 1B) were well-conserved with human sequences. Given this sequence conservation, we used an antibody against human phosphorylated IRE1α to examine the posttranslational activation status of IRE1α in ticks. When treated with UPR inducers thapsigargin and tunicamycin, increased IRE1α phosphorylation was observed in ISE6 tick cells by immunoblot, as expected (Supplemental Figure 2A). *A. phagocytophilum* also induced IRE1α phosphorylation in ISE6 cells, indicating that infection induces receptor activation (Fig 2A). A small molecule inhibitor, KIRA6^60^, successfully blocked IRE1α phosphorylation during infection (Fig 2A) and this inhibition led to significant increases in *A. phagocytophilum* numbers (Fig 2B). Similarly, knocking down the expression of *ire1α*through RNAi also increased *A. phagocytophilum* bacterial burden (Fig 2C). These data show that IRE1α signaling in ticks is activated by infection and restricts bacterial colonization *in vitro*.

**Figure 2.**
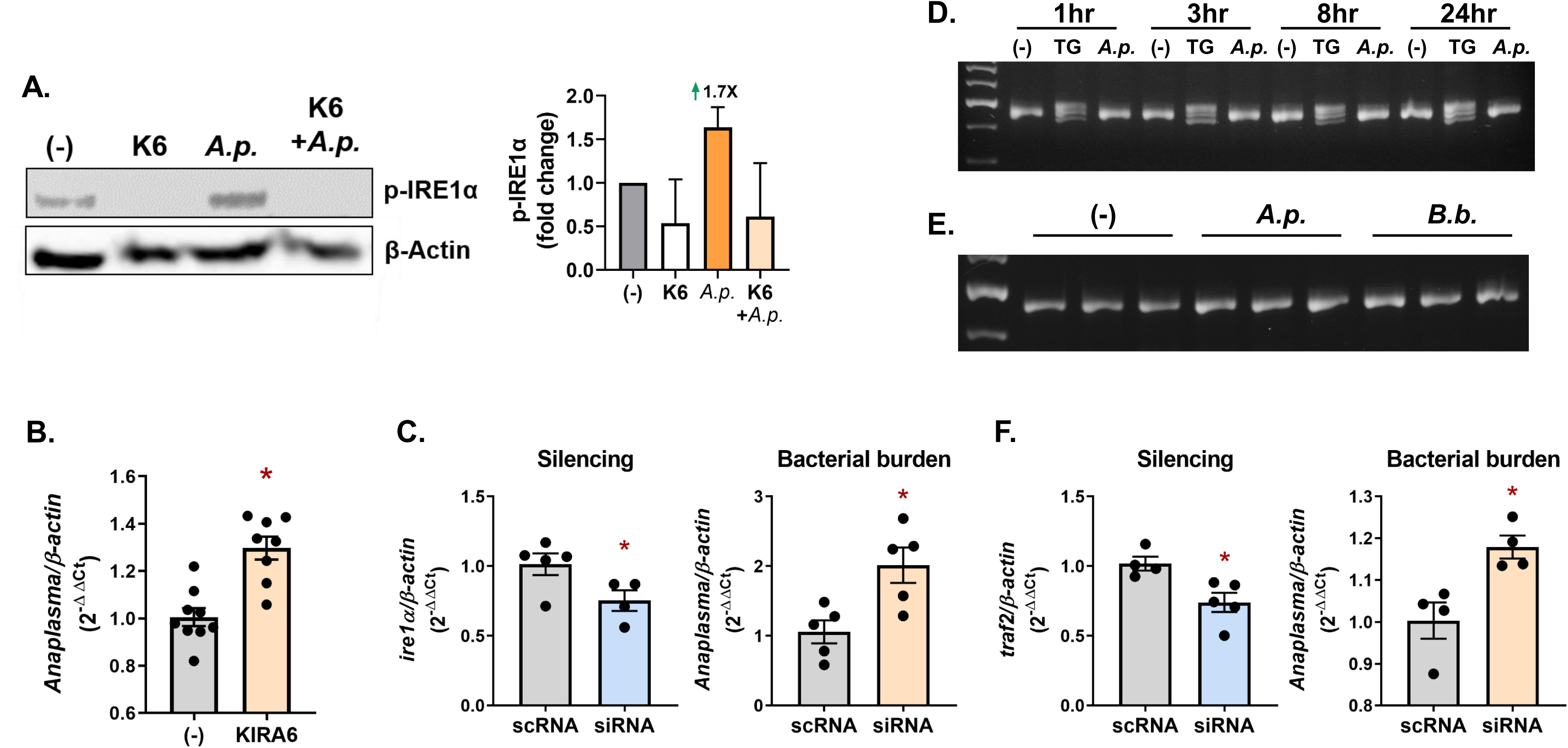
The IRE1α branch of the UPR is induced by *A. phagocytophilum* through TRAF2. **(a)** Phosphorylated IRE1α immunoblot against ISE6 (1×10^6^) cells treated with either the IRE1α inhibitor KIRA6 (K6; 1 hour), infected with *A. phagocytophilum* (*A.p.;* 24 hours) or in combination (1 hour KIRA6 pretreatment, followed by *A. phagocytophilum* infection for 24 hours; K6 + *A.p.*). Immunoblot shown is representative of 2 biological replicates. Protein expression differences were quantified by ImageJ and are expressed as a ratio of phosphorylated IRE1α (∼110 kDa) to the internal loading control, β-actin (45 kDa). ISE6 cells were treated with (**b**) the IRE1α inhibitor KIRA6 (1 hour) or (**c** and **f**) siRNAs to silence gene expression prior to *A. phagocytophilum* (*A.p.*; MOI 50) infection for 18 hours. Gene silencing and *A. phagocytophilum* burden were measured by qRT-PCR. Experiments shown are representative of at least two technical replicates ± SEM. Student’s t-test. *P < 0.05. (**d**) ISE6 cells (1×10^6^) were either untreated (-), stimulated with 0.5 µM of thapsigargin (TG), or infected with *A. phagocytophilum* (*A.p.*; MOI 50) for indicated time points. (**e**) Replete *I. scapularis* nymphs were fed either on uninfected mice (-), *A. phagocytophilum* (*A.p.*)*-*infected, or *B. burgdorferi* (*B.b.*)*-*infected mice. (**d-e**) cDNA was synthesized from RNA and used to evaluate *xbp1* splicing by PCR. Samples were analyzed on a 3% agarose gel. scRNA, scrambled RNA; siRNA, small interfering RNA. See also Supplemental Figure 1 and Supplemental Table 1.

To delineate the signaling events downstream from IRE1α, *xbp1^U^* was next examined in infected ISE6 cells. Primers flanking the *xbp1* intron (Supplemental Fig 1E) were used to differentiate spliced and unspliced transcripts by PCR. Unspliced *xbp1^U^* migrates as a single 459 bp band. In contrast, spliced *xbp1^s^* presents as a trimer on an agarose gel, consisting of spliced transcripts (*xbp1^S^*, 434 bp), unspliced transcripts (*xbp1^U^*) and an *xbp1^U^*-*xbp1^S^* heterodimer that is an artifact of PCR and migrates slightly higher. Spliced *xbp1^s^* was observed in thapsigargin-treated tick cells under all conditions. In contrast, *A. phagocytophilum* infection did not induce *xbp1^U^* splicing at any time points *in vitro* (Fig 2D). We next probed *in vivo* samples from replete *I. scapularis* nymphs that were either fed on uninfected mice or those infected with *A. phagocytophilum* or *B. burgdorferi*. Across all samples, *xbp1^U^* remained unspliced (Fig 2E). These results indicate that although the tick IRE1α is activated by infection and restricts bacterial burden, this phenotype is not carried out through XBP1 activity.

Since XBP1 is not responsive to infection, we sought to determine whether IRE1α is signaling through *Ixodes* TRAF2. Reducing the expression of *traf2* through RNAi in *Ixodes* ISE6 cells caused a significant increase in *A. phagocytophilum* (Fig 2F), correlating with the phenotype observed when silencing *ire1α* transcripts (Fig 2C). These data, together with upregulated *traf2* expression in *A. phagocytophilum*-infected *I. scapularis* nymphs (Fig 1B), suggests that IRE1α is signaling through TRAF2 to restrict pathogen colonization.

### IRE1α interfaces with TRAF2 in I. scapularis ticks

Aligning sequences from humans and ticks reveals that the *Ixodes* TRAF2 is fundamentally unique when compared to the mammalian homolog (Supplemental Figure 3A). The *Ixodes* TRAF2 lacks a RING (Really Interesting New Gene) domain that is necessary for ubiquitin ligase activity^61^. The *Ixodes* TRAF2 also has a reduced TRAF-N domain, which is responsible for bridging interactions with other proteins^132^. Given these differences, we performed homology modeling and a “prediction-driven” docking approach^62^ with the *I. scapularis* IRE1α and TRAF2 proteins to gain insight into how they interact. BLAST was used to identify the human TRAF2 crystal structure^63^ (PDB code 1CA9) as a modeling template for *Ixodes* TRAF2. The modeled form of the *Ixodes* TRAF2 C-terminal region features part of a coiled-coil domain and the highly conserved TRAF-C domain (Supplemental Fig 3B). In addition, the homology model is a trimer where the coiled-coil domain is a single alpha helix and the TRAF-C domain forms an eight-stranded antiparallel β-sandwich. Next, the human IRE1α crystal structure^64^ (PDB code 6URC) was identified by BLAST as a homology template for modeling the cytosolic RNase/kinase domain of *I. scapularis* IRE1α. The structure was modeled in the active state quaternary structure proposed to be necessary for autophosphorylation and RNase activity^65^ (Supplemental Figure 3C-D).

We then modeled the *Ixodes* IRE1α-TRAF2 complex using a prediction-driven docking approach^66^. This tactic combines the utility of interface prediction with *ab initio* docking and is a useful alternative to *ab initio* docking alone when examining protein-protein complex formation. CPORT (Consensus Prediction of interface Residues in Transient complexes)^66^ was used to assign active and passive residues at the interface of the trimeric TRAF-C domains and the RNase/kinase domain of IRE1α (Fig 3A). Residues were then used to filter the docking process by HADDOCK 2.2^67^, which optimizes residue conformations at the interface before proceeding to refinement. The docking model places the trimeric TRAF2 interface at the kinase domain of IRE1α with a buried surface area of 3262.16 Å^2^ (Fig 3B). Importantly, trimeric TRAF2 is positioned in a manner that does not interfere with the IRE1α dimer interface and is away from the C-terminal transmembrane domain (circled) that anchors IRE1α to the ER (Fig 3B). Five salt bridge interactions were identified that define how the TRAF2 trimer is positioned onto the kinase domain of IRE1α (Fig 3C). Each chain of TRAF2 participates in salt bridge interactions with the kinase domain of IRE1α. Therefore, the oligomeric state of TRAF2 seems to play an important role in docking specificity with the RNase/kinase domain of IRE1α. Altogether, *in silico* docking analyses with *Ixodes* IRE1α and TRAF2 suggest that these two molecules can directly interface with one another.

**Figure 3.**
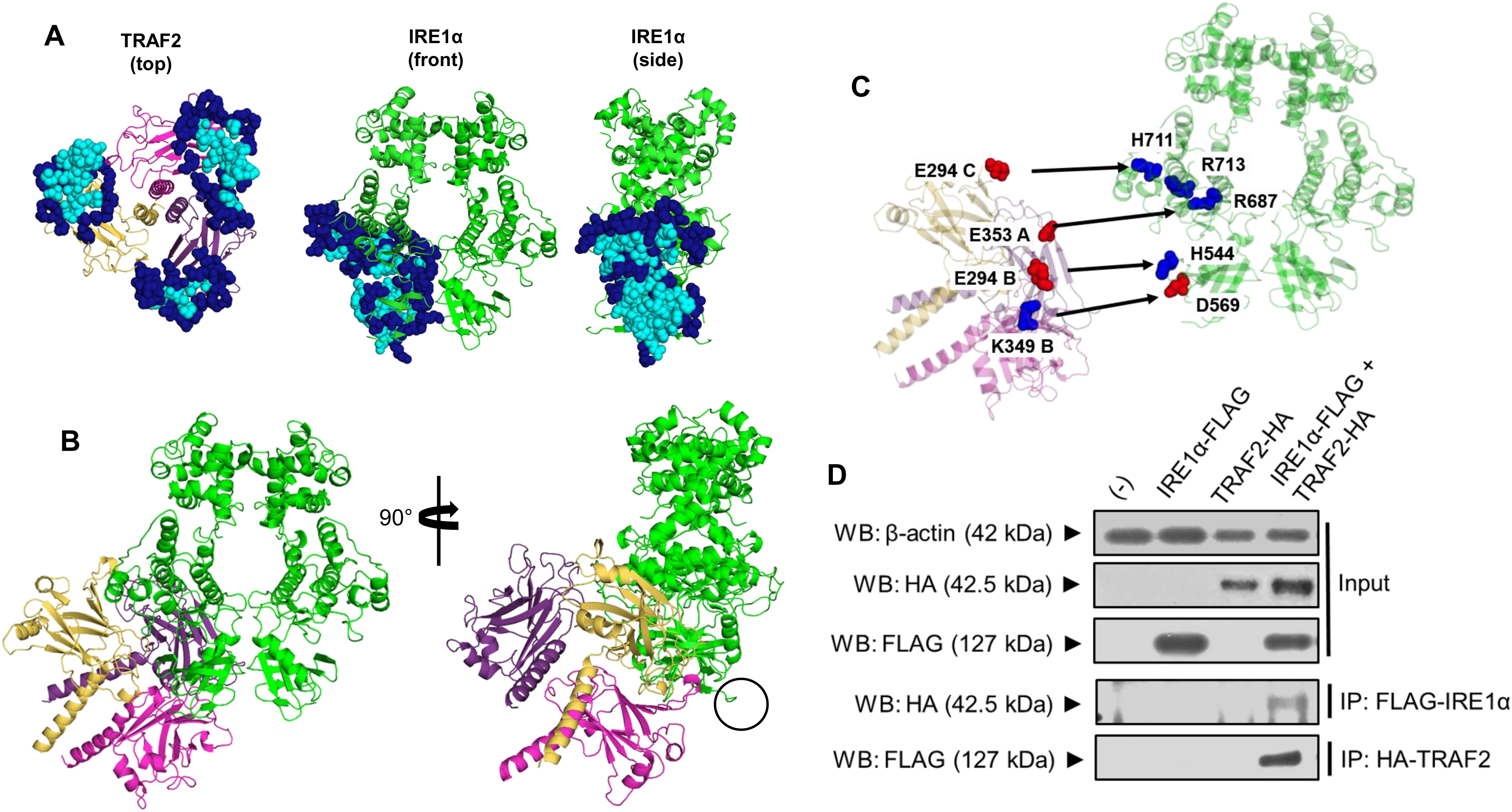
*Ixodes* IRE1α - TRAF2 molecular interactions. (**a**) The interfaces assigned by CPORT for the *Ixodes* TRAF2 trimer and IRE1α homology models. Active central (cyan) and passive peripheral (navy blue) residues shown as spheres were used to filter the docking solutions in HADDOCK 2.2. (**b**) Final docking model between *Ixodes* TRAF2 and IRE1α places TRAF2 away from the dimer interface and the C-terminus of IRE1α (black circle), which anchors IRE1α to the ER. (**c**) Salt bridges were determined between all three chains of *Ixodes* TRAF2 and IRE1α with a measured distance between 2.7 - 2.8 Å. Negatively charged Asp and Glu residues (red spheres) pair with positively charged Lys, Arg, and His residues (blue spheres). (**d**) Immunoprecipitation (IP) analysis followed by Western blotting (WB) showing interaction between FLAG-tagged *Ixodes* IRE1α and HA-tagged *Ixodes* TRAF2 expressed in HEK 293T cells. WB is representative of two biological replicates. See also Supplemental Figure 3 and Supplemental Table 1.

To experimentally validate that IRE1α and TRAF2 specifically interact, we used a Human Embryonic Kidney (HEK) 293T cell transfection system with plasmids expressing *Ixodes* IRE1α and TRAF2 fused to affinity tags (Fig 3D). Recombinant protein expression was confirmed by immunoblotting transfected cells with antibodies for FLAG and HA tags (IRE1α-FLAG and TRAF2-HA). When *Ixodes* IRE1α and TRAF2 are co-expressed, immunoprecipitating with antibodies against the FLAG tag demonstrates that IRE1α specifically pulls down TRAF2 and vice versa (Fig 3D). Altogether, these data demonstrate that *Ixodes* IRE1α and TRAF2 directly and specifically interact.

### Ixodes IRE1α and TRAF2 restrict in vivo bacterial colonization in ticks

We next determined whether the pathogen-restricting activity of *Ixodes* IRE1α and TRAF2 observed *in vitro* had similar impacts *in vivo*. To knock down gene expression, unfed *I. scapularis* nymphs were microinjected with siRNA targeting *ire1α*and *traf2* or with a scrambled control (scRNA). Nymphs were rested overnight and then fed to repletion on *A. phagocytophilum*-infected mice. Gene silencing and bacterial burden were both quantified by qRT-PCR. Similar to *in vitro* experiments, reducing the expression of *ire1α* and *traf2* lead to an increase in *A. phagocytophilum* burdens in *I. scapularis* nymphs (Fig 4A-B).

*I. scapularis* take a blood meal once per life stage, with ticks initially becoming infected during the larval phase^68^. Since gene expression can vary depending on arthropod life stage^69–, 71^, we examined the impact of IRE1α and TRAF2 on pathogen colonization in larvae. We silenced *ire1α* and *traf2* in *I. scapularis* larvae using a modified immersion protocol where ticks were submerged in siRNA or scrambled controls overnight^72^. Following immersion, larvae were rested for 24 hours before feeding to repletion on *A. phagocytophilum*-infected mice. Significant knockdown of *ire1α* and *traf2* was observed in siRNA-treated larvae with this method, which caused an increase in *A. phagocytophilum* numbers (Fig 4C-D).

**Figure 4.**
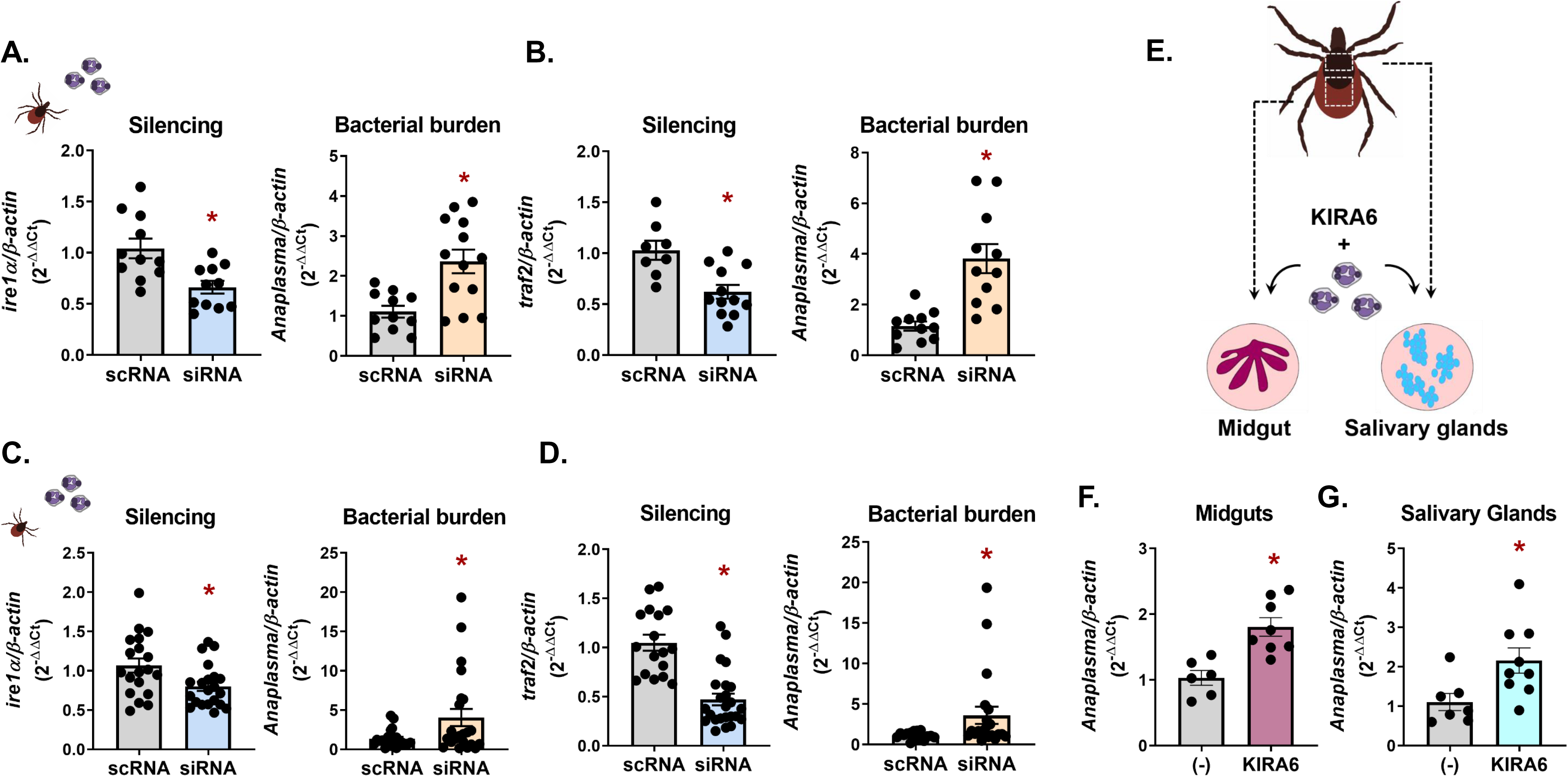
Vector competence for *A. phagocytophilum* is influenced by *Ixodes* IRE1α and TRAF2 at multiple life stages *in vivo.* *I. scapularis* (**a-b**) nymphs or (**c-d**) larvae had *ire1α* and *traf2* expression silenced through RNAi prior to feeding on *A. phagocytophilum-*infected mice. Silencing levels and bacterial load were measured in whole *I. scapularis* nymphs or larvae. (**e**) Schematic of *ex vivo I. scapularis* midgut and salivary gland cultures. (**f-g**) Midguts and salivary glands from *I. scapularis* adults were dissected, cultured, and treated with 1 µM of KIRA6 (1 hour) followed by *A. phagocytophilum* infection for 24 hours. Silencing levels and *A. phagocytophilum* load (16s rDNA) were measured by qRT-PCR. Each point represents 1 tick, midgut, or pair of salivary glands (two technical replicates each), ± SEM. Welch’s t-test. *P < 0.05. scRNA, scrambled RNA; siRNA, small interfering RNA. See also Supplemental Table 1.

Soon after *A. phagocytophilum* is acquired, the bacteria migrate to the salivary glands where they persist throughout the tick life cycle^68, 73, 74^. To understand how IRE1α influences bacterial colonization in tick tissue subsets, we employed an *ex vivo* tick organ culture system^75, 76^. Midguts and salivary glands from adult *I. scapularis* ticks were dissected and treated with the IRE1α inhibitor KIRA6 prior to infection with *A. phagocytophilum* (Fig 5E). Similar to *in vitro* and *in vivo* findings, inhibiting the activity of IRE1α lead to significantly higher *A. phagocytophilum* burdens in *ex vivo* salivary gland and midgut cultures (Fig 4F-G), demonstrating that this signaling axis functionally restricts bacterial colonization in disparate tick tissues.

**Figure 5.**
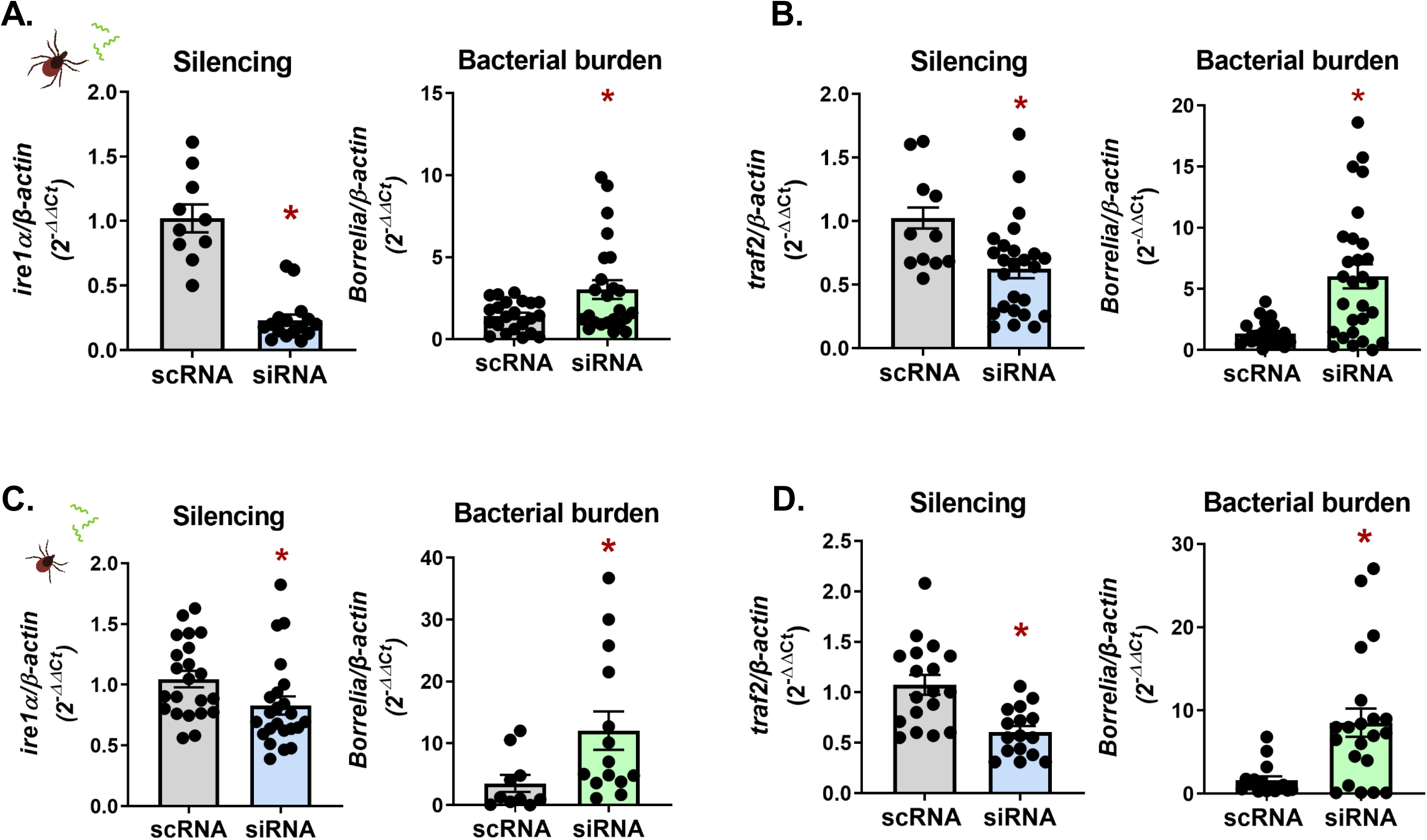
*Ixodes* IRE1α and TRAF2 restrict *B. burgdorferi* colonization *in vivo* at multiple tick life stages. RNAi silencing of *ire1α* and *traf2* in *I. scapularis* (**a-b**) nymphs or (**c-d**) larvae was performed prior to feeding on *B. burgdorferi*-infected mice. Silencing levels and *B. burgdorferi* (*flaB*) were measured in whole *I. scapularis* nymphs or larvae. Each point represents 1 tick (two technical replicates each) ± SEM. Welch’s t-test. *P < 0.05. scRNA, scrambled RNA; siRNA, small interfering RNA. Also see Supplemental Table 1.

We next asked whether the activity of IRE1α-TRAF2 signaling was restrictive to different tick-borne microbes, such as the Lyme disease-causing spirochete *B. burgdorferi*. Expression of *ire1α* and *traf2* was knocked down through RNAi in both *I. scapularis* nymphs and larvae using the same methods described above and ticks were fed to repletion on *B. burgdorferi*-infected mice. In agreement with the phenotype observed with *A. phagocytophilum*, significantly higher *B. burgdorferi* levels were observed in siRNA-treated ticks at both the nymph (Fig. 5A-B) and larval life stages (Fig 5C-D). These data show that IRE1α-TRAF2 signaling is broadly responsive to multiple *I. scapularis*-transmitted pathogens and is functionally restrictive to microbial colonization during different tick life stages.

### The IMD pathway is triggered by IRE1α

TRAF2 is a component of the mammalian TNFR network, which is functionally analogous to the arthropod IMD pathway. This parallel led us to ask whether the antimicrobial activity of the *Ixodes* IRE1α-TRAF2 axis operates through arthropod immunity. AMPs specific to the IMD pathway have not yet been identified in ticks. Instead, the *Drosophila* S2* cell line can be used as a surrogate model to quantify pathway-specific AMPs^28^. To examine whether ER stress induces an immune response in the absence of microbes, we treated S2* cells with the UPR inducer thapsigargin. AMPs corresponding to the IMD pathway (*diptericin*, *attacin A,* and *cecropin A2*^77^) were significantly induced in a dose-dependent manner compared to unstimulated controls (Supplemental Figure 4A). In contrast, the Toll pathway AMP *IM1*^77–79^ was not significantly different, demonstrating that ER stress leads to IMD pathway activation independent of microbial agonists.

It is known that the IMD pathway is responsive to tick-transmitted pathogens *A. phagocytophilum* and *B. burgdorferi*^28, 30^. Since tick-borne microbes also activate the UPR (Figs 1B and 2A) and ER stress induces the IMD network (Supplemental Figure 4A), we asked whether blocking IRE1α during infection would inhibit the IMD pathway. S2* cells that were treated with the IRE1α inhibitor KIRA6 prior to *A. phagocytophilum* or *B. burgdorferi* infection showed significantly reduced IMD pathway AMPs (Supplemental Figure 4B-C).

We next examined whether the tick IMD pathway underwent a similar UPR-driven activation event. Relish is the transcription factor associated with IMD pathway activation. Similar to what was observed in *Drosophila* S2* cells, ISE6 cells that were treated with UPR stimulators thapsigargin or tunicamycin showed an increase in Relish activation (Fig 6A). We next asked if inhibiting IRE1α would block activation of the IMD pathway in ticks. ISE6 cells were stimulated with *A. phagocytophilum* and *B. burgdorferi* alone or were first pretreated with the IRE1α inhibitor, KIRA6, before infection. Pretreatment with KIRA6 resulted in a decline in Relish activation (Fig 6B-C), indicating that infection-induced IMD pathway activation occurs through IRE1α. Collectively, our results provide strong evidence that the IRE1α-TRAF2 axis functions as an IMD pathway-activating mechanism.

**Figure 6.**
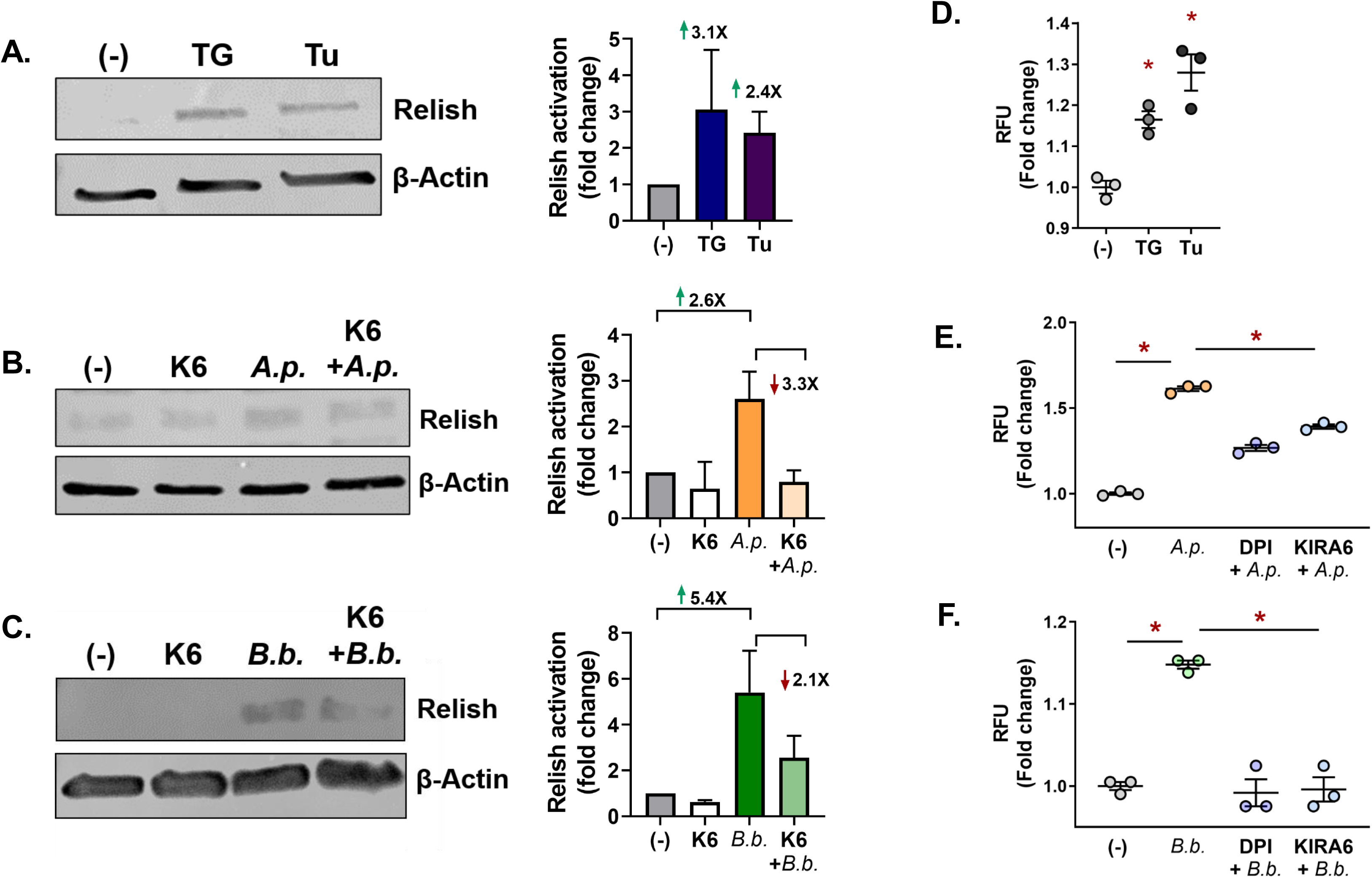
Infection-induced IMD pathway activation and ROS production functions through IRE1α. (**a-c**) Relish immunoblot of ISE6 cells stimulated for 1 hour with (**a**) thapsigargin (TG) or tunicamycin (Tu), or (**b-c**) pretreated with KIRA6 (K6) before (**b**) *A. phagocytophilum* (*A.p.*; MOI 50) or (**c**) *B. burgdorferi* (*B.b.*; MOI 50) infection (24 hours). Immunoblots shown are representative of 2-3 biological replicates. Protein expression differences were quantified by ImageJ and are expressed as a ratio of Relish (∼41 kDa) to the internal loading control, β-actin (45 kDa). (**d-f**) ROS assay with ISE6 cells (1.68×10^5^) stimulated with (**d**) thapsigargin (TG; 10 nM) or tunicamycin (Tu; 50 nM), (**e-f**) ROS output from ISE6 cells pretreated with either DPI (5 µM) or KIRA6 (1 µM) for 1 hour prior to (**e**) *A. phagocytophilum* or (**f**) *B. burgdorferi* infection. ROS was measured by relative fluorescence units (RFU) after 72 hours. Data shown is representative of 3 biological replicates and 2 technical replicates, ± SEM. Student’s t-test. *P < 0.05. (-), vehicle control; DPI, diphenyleneidonium.

### Ixodes IRE1α-TRAF2 signaling potentiates reactive oxygen species

A complementary immune mechanism to the IMD pathway is the production of reactive oxygen species (ROS), which cause bactericidal damage to nucleic acids, proteins, and membrane lipids^15, 80, 81^. Because *B. burgdorferi* and *A. phagocytophilum* are both sensitive to killing by ROS^82–85^ and the mammalian UPR can lead to ROS production^86, 87^, we investigated whether ROS can be induced by the *Ixodes* IRE1α-TRAF2 pathway. ISE6 cells were stimulated with either thapsigargin, tunicamycin, or a vehicle control and monitored for ROS with the fluorescent indicator 2ʹ,7ʹ-dichlorofluorescin diacetate. Pharmacological inducers caused significantly higher fluorescence, indicating that the tick UPR potentiates ROS (Fig 6D).

Infection with *A. phagocytophilum* and *B. burgdorferi* also elicited ROS production in tick cells (Fig 6E-F). Pretreating ISE6 cells with the ROS-inhibiting agent DPI (diphenyleneidonium chloride) prior to infection reduced fluorescence, as expected. Importantly, blocking IRE1α activity with KIRA6 either reduced or completely mitigated ROS (Fig 6E-F), demonstrating that infection-induced ROS is potentiated by IRE1α.

### IRE1α-TRAF2 signaling restricts pathogens across tick vectors

Since the UPR is conserved across eukaryotes, we explored the possibility that the microbe-restricting activity of IRE1α-TRAF2 signaling could functionally impact other arthropod vectors. *D. andersoni* ticks are important disease vectors that transmit several pathogens including the obligate intracellular rickettsia, *A. marginale*^88^. When inducing the UPR in the *D. andersoni* tick cell line DAE100 with tunicamycin and thapsigargin (Fig 7A-B) or blocking IRE1α with KIRA6 (Fig 7C), we observed significant changes in *A. marginale* invasion and replication, comparable to what was observed with *I. scapularis* and *A. phagocytophilum* (Figs 1C-D, 2B). Moreover, higher bacterial loads were also observed in *D. andersoni ex vivo* midgut and salivary gland cultures when IRE1α activity was blocked with KIRA6 (Fig 7D-F). Altogether, this demonstrates that the microbe-restricting activity of IRE1α-TRAF2 signaling is conserved across tick species and is active against disparate pathogens, including intracellular bacteria (*A. phagocytophilum* and *A. marginale*) and extracellular spirochetes (*B. burgdorferi*).

**Figure 7.**
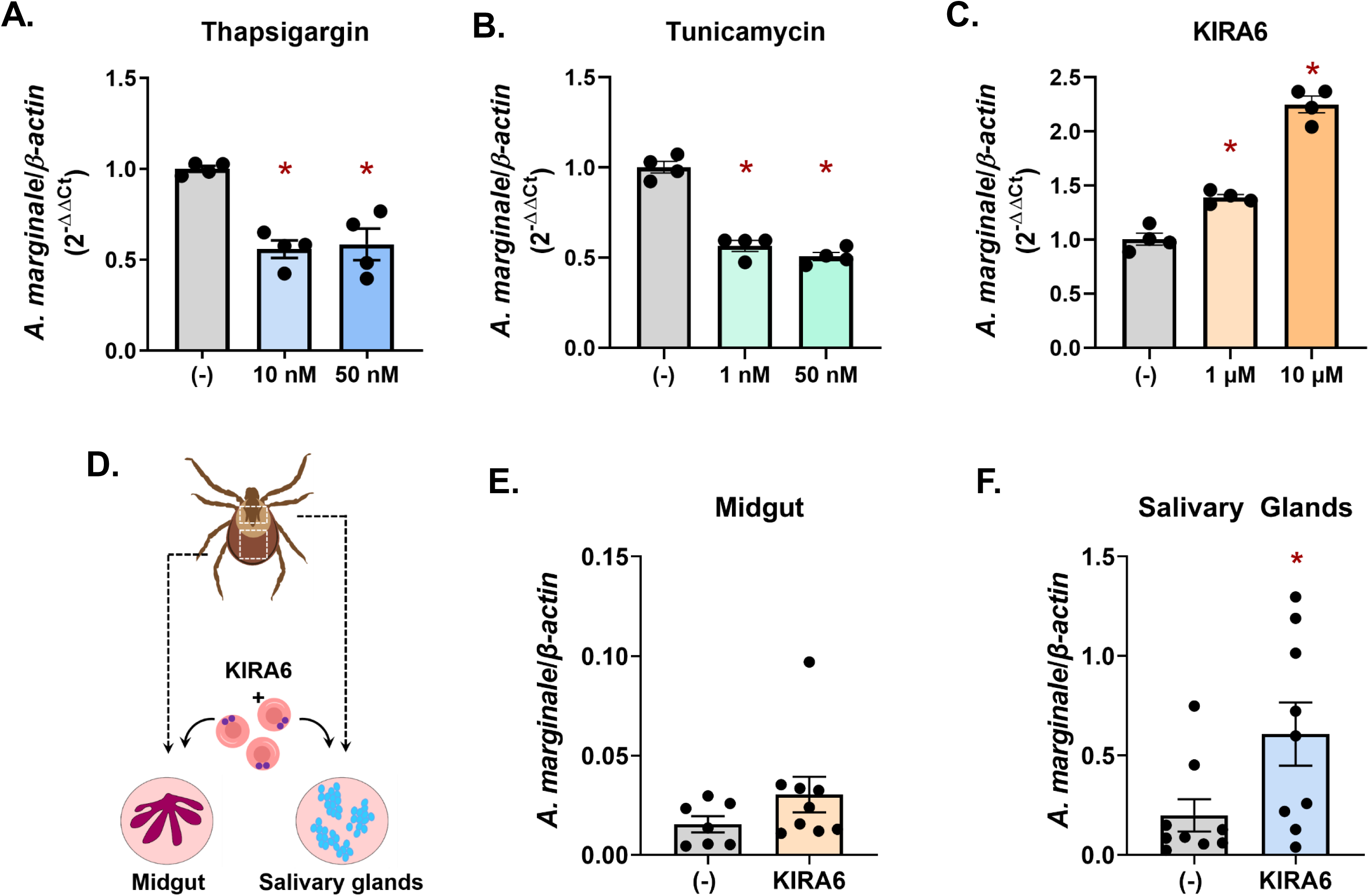
IRE1α and TRAF2-mediated pathogen restriction is conserved across ticks. DAE100 cells (5×10^5^) were treated with indicated concentrations of (**a**) thapsigargin, (**b**) tunicamycin, or (**c**) KIRA6 followed by infection with *A. marginale* (MOI 50) for 18 hours. Student’s t-test. *P < 0.05. (**d**) Schematic of *ex vivo D. andersoni* midgut and salivary gland cultures. (**e-f**) Midguts and salivary glands from *D. andersoni* adults were dissected, cultured, and treated with 1 µM of KIRA6 (1 hour) followed by *A. marginale* infection for 22 hours. *A. marginale* (*rpoH*) was quantified by qRT-PCR and graphed relative to *β-actin.* Welch’s t-test. *P < 0.05. Each point is representative of 1 tick, midgut, or pair of salivary glands (two technical replicates), ± SEM. See also Supplemental Table 1.

## DISCUSSION

How arthropod immunity responds to infection is a fundamental factor influencing the ability of vectors to harbor and transmit pathogens^2–8^. The IMD pathway is increasingly recognized as being divergent across species, with classically defined upstream regulators missing in many arthropod genomes^15–27, 32, 34, 35^. This suggests that an alternative activation mechanism exists. In this article we demonstrate that the *I. scapularis* IMD pathway is initiated through the IRE1α-TRAF2 axis of the UPR. Colonization and replication of *A. phagocytophilum* and *B. burgdorferi* are restricted in ticks by *Ixodes* IRE1α and TRAF2 both *in vitro* and *in vivo*. Moreover, we show that IMD pathway activation and ROS production in response to *A. phagocytophilum* and *B. burgdorferi* are dependent on IRE1α activity and that this mode of antibacterial restriction is conserved across ticks. Collectively, our findings provide an explanation for how the core IMD pathway is activated in the absence of canonical upstream regulators.

To our knowledge, this is the first time that cellular stress responses have been implicated in influencing vector competency. Why host cell stress responses are triggered by *A. phagocytophilum* and *B. burgdorferi* remains unclear. Ticks do not appear to suffer pathological consequences from the microbes they transmit. The connection between host cell stress and immune outcomes supports a model where transmissible pathogens would benefit most by decreasing infection-induced stress. This model is reenforced by the absence of common inflammatory PAMPs in many tick-transmitted pathogens. For example, all *Ixodes*-transmitted bacteria lack lipopolysaccharide (LPS) and DAP-PGN^89–92^. *B. burgdorferi* flagella are housed in the periplasm, effectively shielded from recognition by host cells^93^. During coevolution with ticks, *Ixodes*-transmitted pathogens may have lost inflammatory PAMPs with the benefit of reducing cellular stress and host responses, thereby promoting persistence and transmission. Nevertheless, our data shows that *A. phagocytophilum* and *B. burgdorferi* impart at least some stress on ticks. Since immune responses are energetically costly to the host^94, 95^, we speculate that the tick response is tuned to match the level of threat imposed by infection, ultimately striking a balance that conserves resources and preserves tick fitness.

Our findings indicate a mechanism of IMD pathway activation that deviates from the classically defined paradigm where pattern recognition receptors (PRRs) sense bacterial-derived PAMPs. Both intracellular and extracellular pathogens impart stress on the host, which can be caused by secreted toxic byproducts, competition for nutrients, and/or physical damage to host cells/organ systems^96^. For example, *B. burgdorferi* is an extracellular spirochete and an extreme auxotroph that lacks many central metabolic pathways^97, 98^. To get around this limitation, it parasitizes purines^99^, amino acids^100^, cholesterol^101, 102^, long-chain fatty acids^103, 104^, carbon sources^105^, and other metabolites^106^ from the host. *A. phagocytophilum* is obligately intracellular and parasitizes amino acids and cholesterol from the host, in addition to manipulating host cell processes with secreted effectors^107–112^. From this perspective, both microbes cause stress to the host by competing for a finite amount of resources and disturbing normal cellular processes. Indeed, our evidence shows that tick-transmitted microbes stimulate the UPR and are restricted by its activity. Although cellular stress responses detect and respond to stress, they are not necessarily specific to types of stressors and instead respond by monitoring macromolecular threats to the cell^40, 41, 113^. This more generalized signal widens the infection-sensing scope of possibility and reduces the requirement for an array of specific immune receptors. In this regard, a wide variety of stimuli would converge on a common immune outcome. Since the UPR is an evolutionarily conserved mechanism across eukaryotes^36–38^, it is feasible that UPR-initiated immunity is an ancient mode of pathogen-sensing and host defense against a broad array of infectious organisms.

In summary, we have discovered a linkage between cellular stress responses and arthropod immunity where the *Ixodes* IRE1α-TRAF2 signaling axis initiates the IMD pathway (Supplemental Figure 5). The previous “orphaned” status of the IMD pathway in ticks was a perception that arose from comparative studies with the insect model organism, *Drosophila*. In fact, the absence of upstream IMD pathway regulators appears to be a shared trait among chelicerates and hemimetabolous insects^15–24, 26–30, 32–34, 95^. This revelation underscores the importance of studying fundamental processes outside of model organisms, which may be valuable for determining concepts that could be basally applicable across species. Our findings are conceptually important given that the IMD pathway widely impacts vector competence in many arthropods. With this commonality, one can envision a scenario where a conserved network across species may be an attractive target for future transmission intervention strategies.

## METHODS

### Bacteria and animal models

*E. coli* cultures were grown in lysogeny broth (LB) supplemented with ampicillin at 100 µg µl^-1^. Cultures were grown overnight at 37°C with shaking between 230-250 RPM.

*A. phagocytophilum* strain HZ was cultured in HL60 cells with Roswell Park Memorial Institute (RPMI) 1640 medium supplemented with 10% heat-inactivated fetal bovine serum (Atlanta Biologicals, S11550) and 1X Glutamax (Gibco, 35050061). Cells were maintained between 1×10^5^ - 1×10^6^ ml^-1^ at 37°C, 5% CO_2_. *A. phagocytophilum* was enumerated as previously described^114^. Briefly, the percentage of infected cells is multiplied by the average number of microcolonies per cell, termed ‘morulae’ (5), the average bacteria per morulae (19) and the average amount of bacteria typically recovered from the isolation procedure (50%). Host cell-free *A. phagocytophilum* was isolated by syringe lysis with a 27 gauge needle as previously described^3^.

*B. burgdorferi* B31 (strain MSK5^115^) was grown in modified Barbour-Stoenner-Kelly (BSK) II medium supplemented with 6% normal rabbit serum (NRS, Pel-Freez, 31126-5) at 37°C, 5% CO_2_^115, 116^. Spirochete density and growth phase were monitored by dark field microscopy. Prior to infection, plasmid profiles of all *B. burgdorferi* cultures were screened by PCR, as described previously^115^.

Uninfected *I. scapularis* ticks were provided by the Biodefense and Emerging Infectious Diseases (BEI) Research Resources Repository from the National Institute of Allergy and Infectious Diseases (NIAID) (www.beiresources.org) at the National Institutes of Health (NIH) or from Oklahoma State University (Stillwater, OK, USA). Ticks were maintained in a 23°C incubator with 16/8 hours light/dark photoperiods and 95-100% relative humidity. C3H/HeJ mice were purchased from Jackson Laboratories and C57BL/6 mice were obtained from colonies maintained at Washington State University. 6-10 week old male mice were used for all experiments. C57BL/6 mice were infected intraperitoneally with 1×10^7^ host cell-free *A. phagocytophilum*. C3H/HeJ mice were inoculated intradermally with 1×10^5^ low passage *B. burgdorferi*. All mice were confirmed for infection status prior to tick placement by collecting 25-50 μl of blood from the lateral saphenous vein of each mouse 7 days post-infection. *A. phagocytophilum* burdens were enumerated by quantitative PCR (*16s* relative to mouse *β-actin*^117, 118^)*. B. burgdorferi*-infected blood was subcultured in BSK-II media and examined for the presence of spirochetes by dark field microscopy^119, 120^. Experiments involving mice were carried out according to guidelines and protocols approved by the American Association for Accreditation of Laboratory Animal Care (AAALAC) and by the Office of Campus Veterinarian at Washington State University (Animal Welfare Assurance A3485-01). The animals were housed and maintained in an AAALAC-accredited facility at Washington State University in Pullman, WA. All procedures were approved by the Washington State University Biosafety and Animal Care and Use Committees.

### D. melanogaster and tick cell cultures

*D. melanogaster* S2* cells were cultured with Schneider’s *Drosophila* Medium (Gibco, 21720024) supplemented with 10% heat inactivated FBS (Sigma, SH30070) and 1X Glutamax. Cell were maintained in T75 culture flasks (Corning, 353136) at 28°C.

The *I. scapularis* tick cell line, ISE6, was cultured at 32°C, 1% CO_2_ in L15C-300 medium supplemented with 10% heat inactivated FBS (Sigma, F0926), 10% Tryptose Phosphate Broth (TPB, BD, B260300) and 0.1% Lipoprotein Bovine Cholesterol (LPBC, MP Biomedicals, 219147680)^121^. The *D. andersoni* tick cell line, DAE100, was maintained at 34°C and cultured in L15B medium supplemented with 5% FBS, 10% TBP, and 1% LPBC as previously described^122, 123^.

### Polyacrylamide gel electrophoresis and Western blotting

Protein concentrations were quantified using BCA assays per manufacture protocol (Pierce, 23225). 50 µg of protein per sample were separated on a 4-15% MP TGX precast cassette (Bio-Rad, 4561083) at 100V for 1 hour 25 minutes before being transferred to a PVDF membrane. Membranes were blocked with 5% milk in PBS-T (1X phosphate-buffered saline containing 0.1% Tween-20) for 1-2 hours at room temperature before being incubated at 4°C overnight with a primary antibody in PBS-T with 5% BSA (Bovine Serum Albumin) or 0.5%-5% milk. Primary antibodies used for immunoblotting are as follows: α-phospho-IRE1α (Abcam, ab124945, 1:1000), α-Relish (gift from Joao Pedra; 1:500), α-Actin (Sigma, A2103, 1:1000), α-HA (Pierce, 26183, 1:1000), and α-FLAG-HRP (Sigma, A8592, 1:500). Secondary antibodies were applied for 1-2 hours at room temperature and are as follows: Goat α-Rabbit-HRP (Abcam, ab97051, 1:5000), Donkey α-Rabbit-HRP (Thermo Fisher Scientific, A16023, 1:2000), Rabbit α-Mouse-HRP (Bio-Rad, STAR13B, 1:2000), and Rec-G-Protein-HRP (Thermo Fisher Scientific, 101223, 1:2000). Blots were visualized with Enhanced Chemiluminescence (ECL) Western blotting substrate (Thermo Fisher Scientific, 32106). If necessary, blots were stripped with Western Blot Stripping Buffer (Thermo Fisher Scientific, 21059) for 15-20 minutes at room temperature with shaking.

### Plasmid construction

Both *Ixodes* IRE1α and TRAF2 were codon optimized for expression in human cell lines (GenScript). Primers listed in Supplemental Table 1 were used to amplify full length *I. scapularis ire1α* for cloning into pCMV/hygro-Negative Control Vector (SinoBiological, CV005) with *HindIII* sites. Full length *I. scapularis traf2* was amplified and cloned into pCMV-HA (New MCS) vector (received as a gift from Christopher A. Walsh; Addgene plasmid #32530) using *XhoI* and *EcoRV*. All constructs were confirmed by sequencing (Eurofins Genomics).

### Maintenance and Transfection of HEK 293T cells

HEK 293T cells were cultured in Dulbecco’s Modified Eagle’s Medium (DMEM, Sigma, D6429) supplemented with 10% heat inactivated FBS (Atlanta Biologicals, S11550) and 1X Glutamax. Cells were maintained in T75 culture flasks (Corning, 353136) at 37°C, 5% CO_2_. For transfection, 1×10^6^ HEK 293T cells were seeded into 6-well plates and allowed to attach overnight. The following day cells were transfected with 2.5 µg of pCMV-TRAF2-HA and/or pCMV-IRE1α-FLAG plasmid DNA using 10 µl of Lipofectamine 2,000 (Invitrogen, 11668027) in Opti-MEM I Reduced Serum Medium (Gibco, 31985062). After 5 hours, media containing the plasmid-Lipofectamine 2,000 complex was removed and replaced with complete DMEM for 48 hours at 33°C, 5% CO_2_. The transfected cells were lysed with 500 µl of 25 mM Tris-HCl pH 7.4, 150 mM NaCl, 1% NP-40, 1 mM EDTA, 5% glycerol with 1X protease and phosphatase inhibitor cocktail (Thermo Scientific, 78440) for 15 minutes on ice.

### Co-immunoprecipitation assay

*Ixodes* IRE1α-FLAG and TRAF2-HA expression was validated by immunoblotting whole cell lysates with α-FLAG-HRP (Sigma, A8592, 1:500) and α-HA (Pierce, 26183, 1:1000). After protein expression was confirmed, cross-linked agarose beads (α-FLAG M2: Sigma, A2220; α-HA: Pierce, 26181) were washed 2X with TBS (50 mM Tris, 150 mM NaCl, pH 7.5) and incubated with lysis buffer at 4°C for 1 hour. Approximately 1-2 mg of cell lysate was combined with 80 µl (packed volume) of cross-linked agarose beads and incubated overnight at 4°C. Beads were washed 3 times with TBS and protein was eluted by boiling in 50 µl of 4X Laemmli buffer for 5 minutes. Protein interactions were evaluated by immunoblot as described above.

### Template-based homology modeling of Ixodes TRAF2 and the RNase/kinase domain of IRE1α

A BLAST search in the Protein Data Bank (PDB) using the *Ixodes* TRAF2 sequence returned the candidate template crystal structure of the TRAF-C domain from human TRAF2 (39.64% sequence identity). The human TRAF2 crystal structure (PDB code 1CA9) was used as a reference for building the homology model of the TRAF-C domain and part of the coiled-coil domain for *Ixodes* TRAF2 (residues 176-357) in SWISS-MODEL^63, 124^. QMEANdisCo was used to obtain a quality score, which defines how well the homology model aligns to reference structures in the PDB. Scores closer to 1 indicate that the homology model matches well to other reference structures^125^. Quality assessment of the TRAF2 homology model in QMEANDisCo gave a score of 0.69. The GalaxyRefine server was used to then further refine the *Ixodes* TRAF2 homology model, which increased the quality score in QMEANDisCo to 0.71^126^.

A PDB BLAST search for *Ixodes* IRE1α returned the candidate template crystal structure of the RNase/kinase domain from human IRE1α (62.20% sequence identity). A homology model for the cytosolic RNase/kinase domain of tick IRE1α (residues 525-944) was built using the crystal structure of the RNase/kinase domain from humans (PDB code 6URC) with SWISS-MODEL^64, 124^. Quality assessment of the tick IRE1α homology model in QMEANDisCo gave a score of 0.78.

### Prediction-driven docking of Ixodes TRAF2 and the RNase/kinase domain of IRE1α

A consensus interface predictor, CPORT (Consensus Prediction of interface Residues in Transient complexes), was used to assign residues at the interface of *Ixodes* TRAF2 and the IRE1α RNase/kinase domain^66^. Predicted residues were used to define the docking interface between *Ixodes* TRAF2 and IRE1α for docking in HADDOCK2.2^67^. The docked model was immersed in a solvent shell using the TIP3P water model and a short 300K MD simulation was ran to optimize side chains and improve interaction energetics^67^. The cluster with the lowest Z-score was chosen for further analysis. Docking models were then screened based on salt bridge interactions at the docking interface and the model with the best chemical complementarity was used in the final analysis. PyMOL version 2.2.3 was used for all distance measurements of salt-bridge interactions (<4 Å cutoff) (The PyMOL Molecular Graphics System, Schrodinger, LLC).

### ROS assay

ISE6 cells were seeded at a density of 1.68×10^5^ cells per well in a black-walled, clear-bottom 96-well plate (Thermo Scientific, 165305) with L15C-300 media. The cells were maintained in growth conditions described above for the length of experiments. All wells were treated for 1 hour with 10 µM 2’,7’-dichlorofluorescin diacetate (DCF-DA, Sigma, D6883) in Ringer buffer (155 mM NaCl, 5 mM KCl, 1 mM MgCl_2_ · 6H_2_O, 2 mM NaH_2_PO_4_ · H_2_O, 10 mM HEPES, and 10 mM glucose)^127^ alone or with 5 µM diphenyleneidonium chloride (DPI, Sigma, D2926), 1 µM KIRA6 (Cayman Chemical, 19151), or 0.1% DMSO. Buffer was removed; cells were washed with room temperature 1X PBS and incubated for 72 hours in L15C-300 alone or with *A. phagocytophilum* (MOI 200), *B. burgdorferi* (MOI 200), 10 nM thapsigargin (TG, Sigma, T9033), or 50 nM tunicamycin (Tu, Sigma T7765). Fluorescence was measured at 504 nm (excitation), 529 nm (emission). Data is graphed as fold change of relative fluorescence units (RFU) normalized to the negative control ± standard errors of the means (SEM).

### Pharmacological treatments, RNAi silencing, quantitative reverse transcriptase-PCR

ISE6 cells were seeded at 1×10^6^ cells per well and DAE100 cells were seeded at 5×10^5^ cells per well in a 24-well plate and pre-treated with KIRA6, thapsigargin, or tunicamycin for indicated times and concentrations prior to infection. Cells were infected with *A. phagocytophilum* (ISE6) or *A. marginale* (DAE100) at an MOI 50 for 18 hours before collection in Trizol (Invitrogen, 15596026). For DAE100 experiments, all incubations occurred at 34°C in a BD campy bag with no gaspak. RNA was extracted using the Direct-zol RNA microprep Kit (Zymo, R2062). cDNA was synthesized from 300-500 ng total RNA with the Verso cDNA Synthesis Kit (Thermo Fisher Scientific, AB1453B). Bacterial burden and gene silencing were assessed by quantitative reverse transcription-PCR (qRT-PCR) with the iTaq Universal SYBR Green Supermix (Bio-Rad, 1725125) using primers listed in Supplemental Table 1. Cycle conditions are as recommended by the manufacturer.

For transfection experiments, siRNAs and scrambled controls (scRNAs) were synthesized following directions from the Silencer siRNA Construction Kit (Invitrogen, AM1620) using the primers listed in Supplemental Table 1. siRNA or scRNA (3 µg) was used to transfect 1×10^6^ ISE6 cells overnight with 2.5 µl of Lipofectamine 2,000. Cells were infected with *A. phagocytophilum* (MOI 50) for 18 hours before being collected in Trizol. RNA was isolated and transcripts were quantified by qRT-PCR as described above. All data are expressed as means ± SEM.

### Ex vivo I. scapularis and D. andersoni organ culture

Ten male and female unfed adult *I. scapularis* ticks were surface sterilized with continuous agitation in 10% benzalkonium chloride (Sigma, 12060) for 10 minutes, washed twice with sterile water, dried on sterile filter paper under aseptic conditions, and transferred to a sterile tube. Midgut and salivary glands were excised on a microscope slide in a pool of sterile 1X PBS with 100 I.U ml^-1^ penicillin and 100 µg ml^-1^ streptomycin (Gibco, 15140122). Tissues were placed in individual wells of a 96-well plate (Costar, 3595) with 100 µl of L15C-300 and incubated at 32°C with 1% CO_2_. Tissues were treated with 1 µM of KIRA6 or 1% DMSO for 1 hour before the addition of 1×10^6^ *A. phagocytophilum*. 24 hours post-infection, samples were collected following the addition of 100 µl of Trizol. Tissues were homogenized using QIAshredder columns (Qiagen, 79654) according to the manufacturer’s instructions prior to RNA extraction and qRT-PCR analysis, performed as previously described.

Twenty male unfed adult *D. andersoni* ticks were surface sterilized and dissected as above. Tissues were placed in individual wells of a 96-well plate with 100 µl of L15B. Tissues were pretreated with KIRA6 or vehicle control (DMSO) as previously stated prior to the addition of 1×10^6^ *A. marginale* for 22 hours. Samples were collected and processed as above with qRT-PCR standard curves using primers listed in Supplemental Table 1. All data are expressed as means ± SEM

### RNAi silencing in nymphs and larvae

*I. scapularis* nymphs were microinjected as described previously^114, 121^. 10 µl Drummond microdispensers (DrummondSci, 3000203G/X) were drawn to fine point needles using a Narishige PC-100 micropipette puller. *I. scapularis* nymphs were microinjected with 25 nl of siRNA or scRNA (∼1000 ng/µl) into the anal pore using a Dummond Nanoject III Nanoliter Injector (DrummondSci, 3000207). Ticks were allowed to rest overnight before being placed between the ears and on the back of an infected mouse. Each group was placed on a single mouse and fed to repletion (5-7 days). Nymphs were flash frozen with liquid nitrogen, individually crushed with a plastic pestle and suspended in Trizol for RNA extraction.

*I. scapularis* larvae were pre-chilled at 4°C for 5 minutes. Approximately 150 larvae were transferred to a 1.5 ml tube with 40-50 µl of either siRNA or scRNA (∼1000 ng/µl). The tubes were centrifuged at 3000 x g for 5 minutes to encourage submersion of the larvae in the dsRNA and were then incubated overnight at 15°C. The following day, ticks were dried and rested overnight before being placed on mice to feed until repletion (3-7 days). Larvae were flash frozen in liquid nitrogen and individually crushed with a plastic pestle. Trizol was added before proceeding to RNA isolation and qRT-PCR analysis, performed as previously stated.

### xbp1 PCR and agarose gel electrophoresis

RNA was isolated from both ISE6 cells or replete *I. scapularis* nymphs (uninfected, *A. phagocytophilum-*infected, or *B. burgdorferi-*infected). ISE6 cells were treated with either 0.5 µM thapsigargin or *A. phagocytophilum* at an MOI of 50. Cells were collected 1, 3, 8, and 24 hours post-treatment in Trizol. RNA was isolated and cDNA synthesized as previously described. The cleavage status of *xbp1* was assessed via PCR using DreamTaq Green PCR Mastermix (Thermo Scientific, K1082) and the *xbp1* primers listed in Supplemental Table 1 with the cycling protocol recommended by the manufacturer. Samples were analyzed using a 3% agarose (Thermo Fisher, BP160) gel in 1X Tris-Borate EDTA (TBE, Thermo Fisher, BP1333) with 0.5 µg ml^-1^ of ethidium bromide (Thermo Fisher, BP102) and imaged with a Protein Simple AlphaImager HP system.

### UPR and IMD Gene Expression Profiling

Untreated *I. scapularis* nymphs were fed to repletion on *A. phagocytophilum*-infected mice or uninfected mice and frozen. The expression levels of UPR genes were assessed in individual ticks by qRT-PCR as previously described. Primers specific for *bip*, *ire1α*, *xbp1*, and *traf2* are listed in Supplemental Table 1. Data are expressed as means ± SEM.

1×10^6^ *D. melanogaster* S2* cells were seeded in Schneider’s media with 1 µM of 20-hydroxyecdysone to prime the IMD pathway, as previous reported^128^. Cells were treated with indicated concentrations of thapsigargin for 6 hours or with 10 µM of KIRA6 for 1 hour prior to infection with *A. phagocytophilum* (MOI 50) or *B. burgdorferi* (MOI 50) for 6 hours. Samples were collected in Trizol and RNA was isolated. IMD pathway and Toll pathway-specific AMPs were quantified by qRT-PCR with primers listed in Supplemental Table 1 as previously described.

### Gene alignment

UPR gene sequences were identified by querying the *I. scapularis* genome with *Homo sapiens* protein sequences using NCBI (National Center for Biotechnology Information) protein BLAST. Human sequences include BiP (NP_005338.1), IRE1α (NP_001424.3), TRAF2 (NP_066961.2), and XBP1 (NP_005071.2). Human and tick sequences were aligned using Jal view^129^. Shaded regions indicate amino acid physiochemical property conservation. EMBL-EBI (European Bioinformatics Institute) Pfam 34.0 was used to identify and annotate protein domains^130^.

### Statistical analysis

*In vitro* experiments were performed with 3-5 replicates. *In vivo* experiments involved the use of 10-20 ticks. Data were expressed as means ± SEM and analyzed with either unpaired Student’s t*-*test or Welch’s t*-*test. Calculations and graphs were created with GraphPad Prism version 9.0. P *<* 0.05 was considered statistically significant.

## Supporting information

Sidak-Loftis, L.C. et al Supplementary materials

---

**Supplemental Figure 1 (Related to Figure 2) UPR molecules are conserved in the *I. scapularis* genome.** (**a-d**) Amino acid sequence alignment of BiP, IRE1α, TRAF2, and XBP1 between *H. sapiens* and *I. scapularis.* Alignments were created using available sequences from NCBI imported into Jal view. Shaded regions indicate amino acid physiochemical property conservation. Good conservation between sequences was observed for (**a**) BiP, (**b**) the IRE1α protein kinase domain (light blue box) and RNAse domain (grey box), (**c**) the TRAF-type zinc finger domain of TRAF2 (yellow box), and (**d**) the basic region leucine zipper (bZIP) domain in XBP1 (blue box). (**e**) Nucleotide sequence for *I. scapularis xbp1* mRNA. The internal intron that is spliced by the RNase domain of IRE1α is underlined in blue. Cleavage sites are indicated by red lettering and black arrows indicate primer sites used to confirm *xbp1* splicing by PCR.

**Supplemental Figure 2 (Related to Figure 2) Thapsigargin and tunicamycin induce IRE1α phosphorylation in tick cells.** (**a**) Phosphorylated IRE1α immunoblot against ISE6 (1×10^6^) cells treated with ER stress inducers tunicamycin (Tu; 50 nM) and thapsigargin (TG; 50 nM) for 24 hours. Immunoblot shown is representative of 2 biological replicates. Protein expression differences were quantified by ImageJ and are expressed as a ratio of phosphorylated IRE1α (∼110 kDa) to the internal loading control, β-actin (45 kDa).

**Supplemental Figure 3 (Related to Figure 3) *Ixodes* IRE1α and TRAF2 homology models.** (a) Domain comparison between human TRAF2 and *Ixodes* TRAF2 proteins. Really Interesting New Gene (RING; orange). Zinc finger (green). TRAF N-domain (blue). TRAF C domain (red). (b) The *Ixodes* TRAF2 homology model is a trimer with three chains labeled A (purple), B (magenta), and C (yellow). Part of the coiled-coil domain is modeled as a single alpha helix and the TRAF-C domain forms an eight-stranded antiparallel β-sandwich. (**c**) *Ixodes* IRE1α homology model of the dimer RNase/kinase domain. The kinase region consists of an N-lobe (yellow) and C-lobe (aqua). (**d**) Residues of the kinase-extension nuclease (KEN) domain (top) and the kinase domain (bottom) are predicted to participate in salt bridge formation and dimerization. Residues at the KEN domain interface predicted to form salt bridges are shown in the top panel. Residues at the nucleotide binding pocket coordinate MgADP and are conserved with human IRE1α (middle panel). Salt bridge-forming residues at the kinase domain are predicted to participate in IRE1α dimerization (bottom panel).

**Supplemental Figure 4 (Related to Figure 6) The UPR stimulates IMD pathway-associated antimicrobial peptides.** (**a**) Indicated concentrations of thapsigargin (TG) were used to treat S2* cells (1×10^6^) for 6 hours prior to examining gene expression differences. (**b-c**) S2* cells (1×10^6^) were pretreated with KIRA6 (1 hour) before (**b**) *A. phagocytophilum* (*A.p.*; MOI 50) or (**c**) *B. burgdorferi* (*B.b.*; MOI 50) infection (6 hours). Gene expression is relative to *rp49.* Dotted line denotes unstimulated controls. Data shown are representative of 4-5 biological replicates and two technical replicates. See also Supplemental Table 1.

**Supplemental Figure 5 The UPR triggers the IMD pathway in ticks.** Tick-borne bacteria *A. phagocytophilum* and *B. burgdorferi* stimulate the UPR in *I. scapularis* ticks. IRE1α is activated by phosphorylation (P) and pairs with TRAF2. This signaling axis induces the IMD pathway, Relish activation, and antimicrobial responses that restrict pathogen colonization.

## Acknowledgements

We are grateful to Ulrike Munderloh (University of Minnesota) for providing ISE6 and DAE100 tick cell lines; Jon Skare (Texas A&M Health Science Center) for providing *B. burgdorferi* B31 (MSK5); BEI Resources and Oklahoma State University for *Ixodes scapularis* ticks, and for the Addgene plasmid #32530 which was received as a gift from Christopher A. Walsh.

## Author contributions

L.C.S-L., K.L.R. and D.K.S. designed the study. L.C.S-L., K.L.R., N.P., J.K.U, J.H., and D.K.S. performed experiments. N.P. and J.W.P. performed homology modeling and IRE1α-TRAF2 docking. S.M.N. and J.K.U. contributed reagents and performed *D. andersoni* experiments. E.A.F. designed and constructed Supplemental Figure 5. L.C.S-L, K.L.R., N.P., A.S.G, J.W.P., and D.K.S. analyzed data. All authors provided intellectual input into the study. L.C.S-L., K.L.R., N.P., and D.K.S. wrote the manuscript; all authors contributed to editing.

