## Supplementary material for "The unfolded protein response triggers the immune deficiency pathway in ticks": Sidak-Loftis, L.C. et al Supplementary materials

### Slide 1
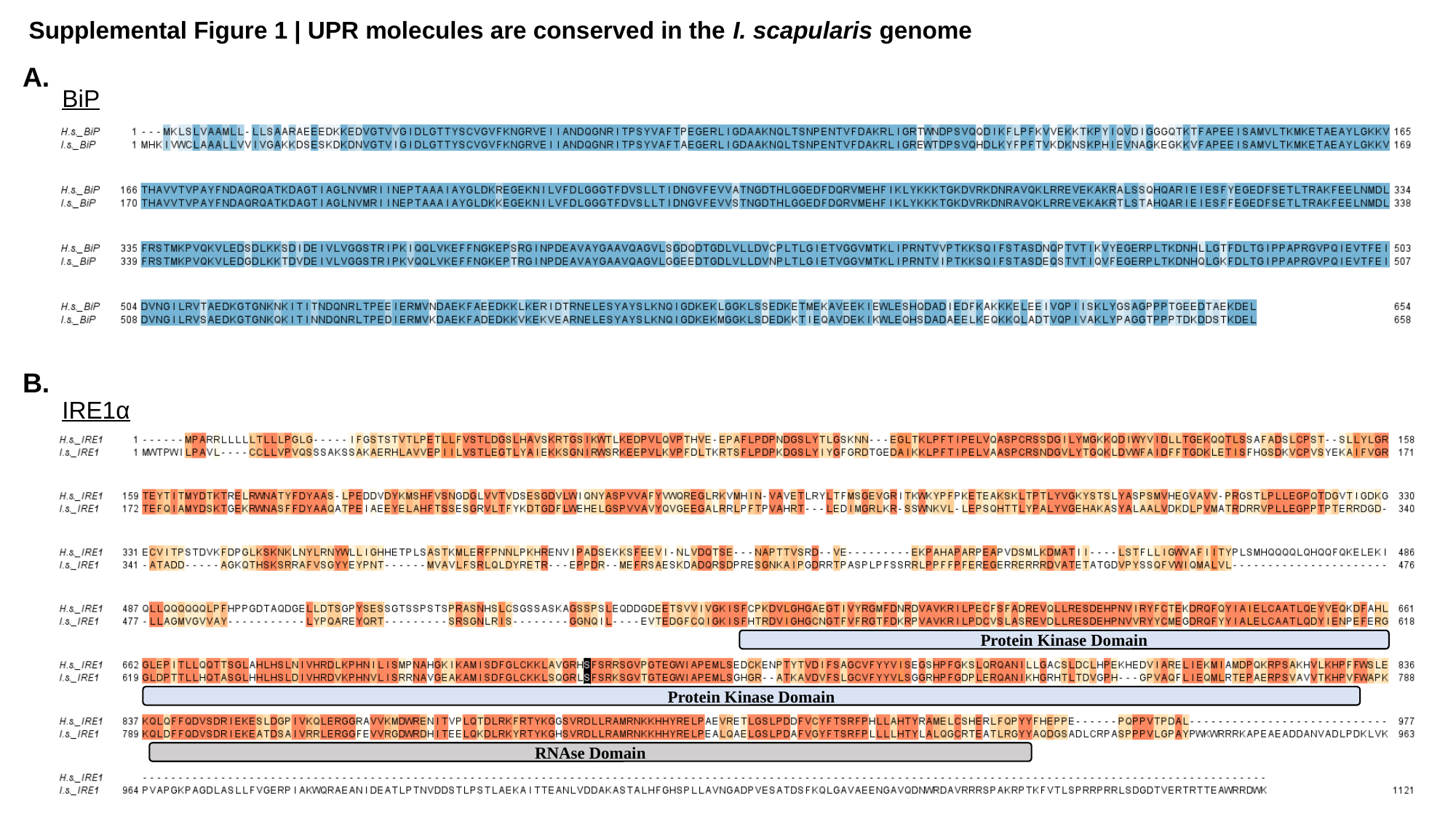

Supplemental Figure 1 | UPR molecules are conserved in the I. scapularis genome
A.
BiP
B.
IRE1α
Protein Kinase Domain
Protein Kinase Domain
RNAse Domain

### Slide 2
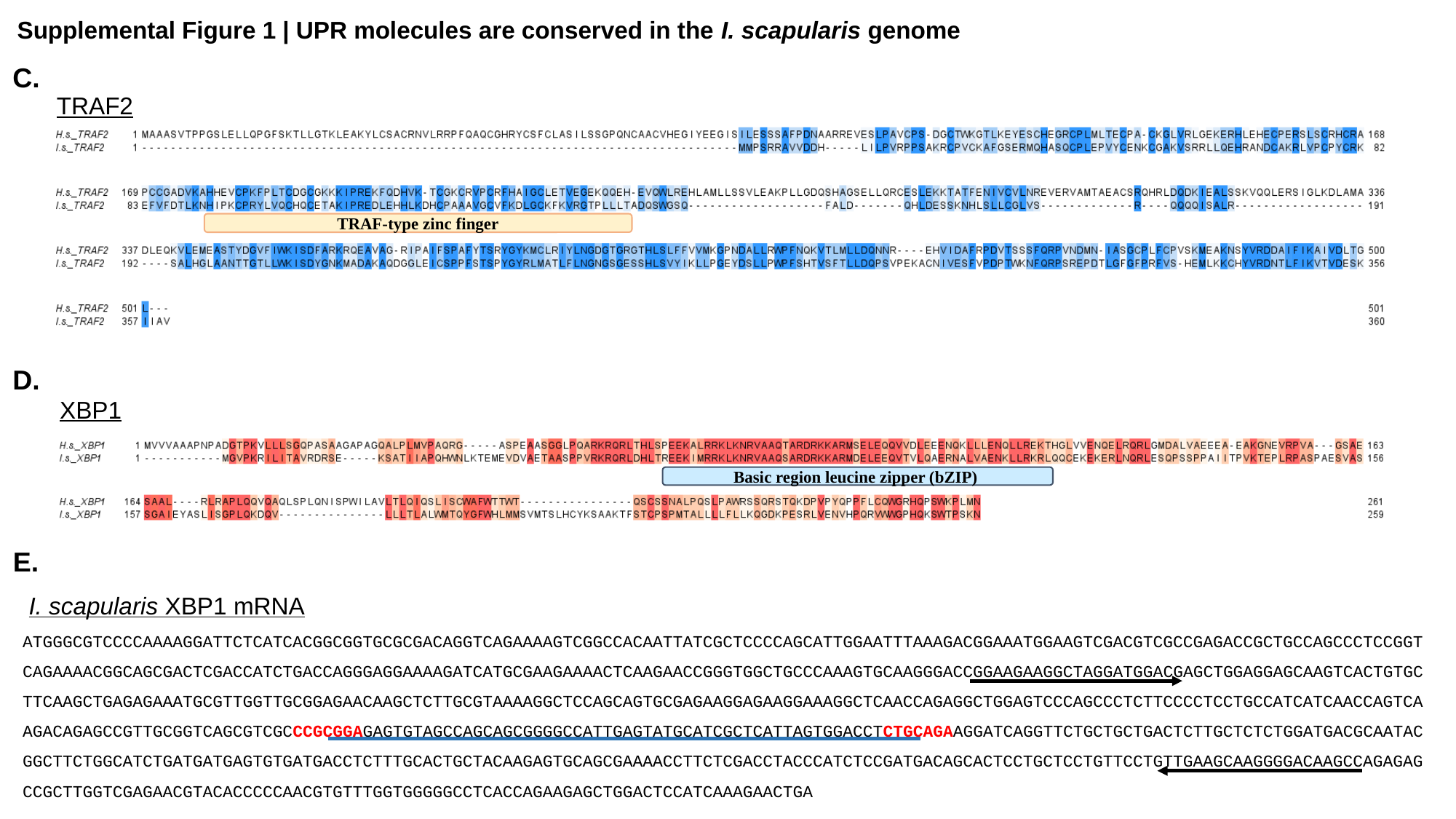

Supplemental Figure 1 | UPR molecules are conserved in the I. scapularis genome
C.
TRAF2
TRAF-type zinc finger
D.
XBP1
Basic region leucine zipper (bZIP)
E.
I. scapularis XBP1 mRNA
atGGGCGtCCCCaaaaGGattCtCatCaCGGCGGtGCGCGaCaGGtCaGaaaaGtCGGCCaCaattatCGCtCCCCaGCattGGaatttaaaGaCGGaaatGGaaGtCGaCGtCGCCGaGaCCGCtGCCaGCCCtCCGGtCaGaaaaCGGCaGCGaCtCGaCCatCtGaCCaGGGaGGaaaaGatCatGCGaaGaaaaCtCaaGaaCCGGGtGGCtGCCCaaaGtGCaaGGGaCCGGaaGaaGGCtaGGatGGaCGaGCtGGaGGaGCaaGtCaCtGtGCttCaaGCtGaGaGaaatGCGttGGttGCGGaGaaCaaGCtCttGCGtaaaaGGCtCCaGCaGtGCGaGaaGGaGaaGGaaaGGCtCaaCCaGaGGCtGGaGtCCCaGCCCtCttCCCCtCCtGCCatCatCaaCCaGtCaaGaCaGaGCCGttGCGGtCaGCGtCGCCCGCGGaGaGtGtaGCCaGCaGCGGGGCCattGaGtatGCatCGCtCattaGtGGaCCtCtGCaGaaGGatCaGGttCtGCtGCtGaCtCttGCtCtCtGGatGaCGCaataCGGCttCtGGCatCtGatGatGaGtGtGatGaCCtCtttGCaCtGCtaCaaGaGtGCaGCGaaaaCCttCtCGaCCtaCCCatCtCCGatGaCaGCaCtCCtGCtCCtGttCCtGttGaaGCaaGGGGaCaaGCCaGaGaGCCGCttGGtCGaGaaCGtaCaCCCCCaaCGtGtttGGtGGGGGCCtCaCCaGaaGaGCtGGaCtCCatCaaaGaaCtGa

### Slide 3
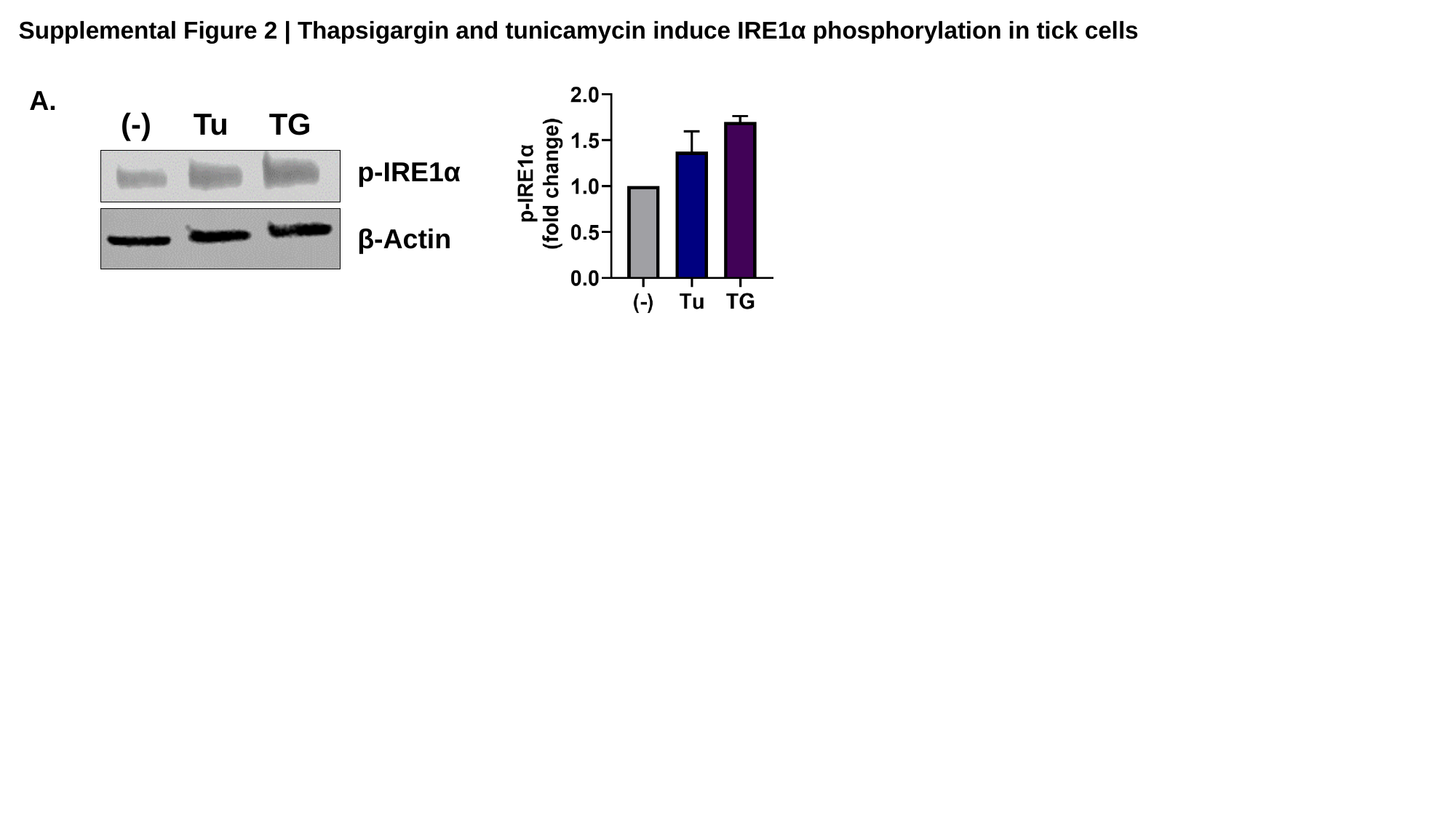

Supplemental Figure 2 | Thapsigargin and tunicamycin induce IRE1α phosphorylation in tick cells
A.
(-)
Tu
TG
p-IRE1α
β-Actin

### Slide 4
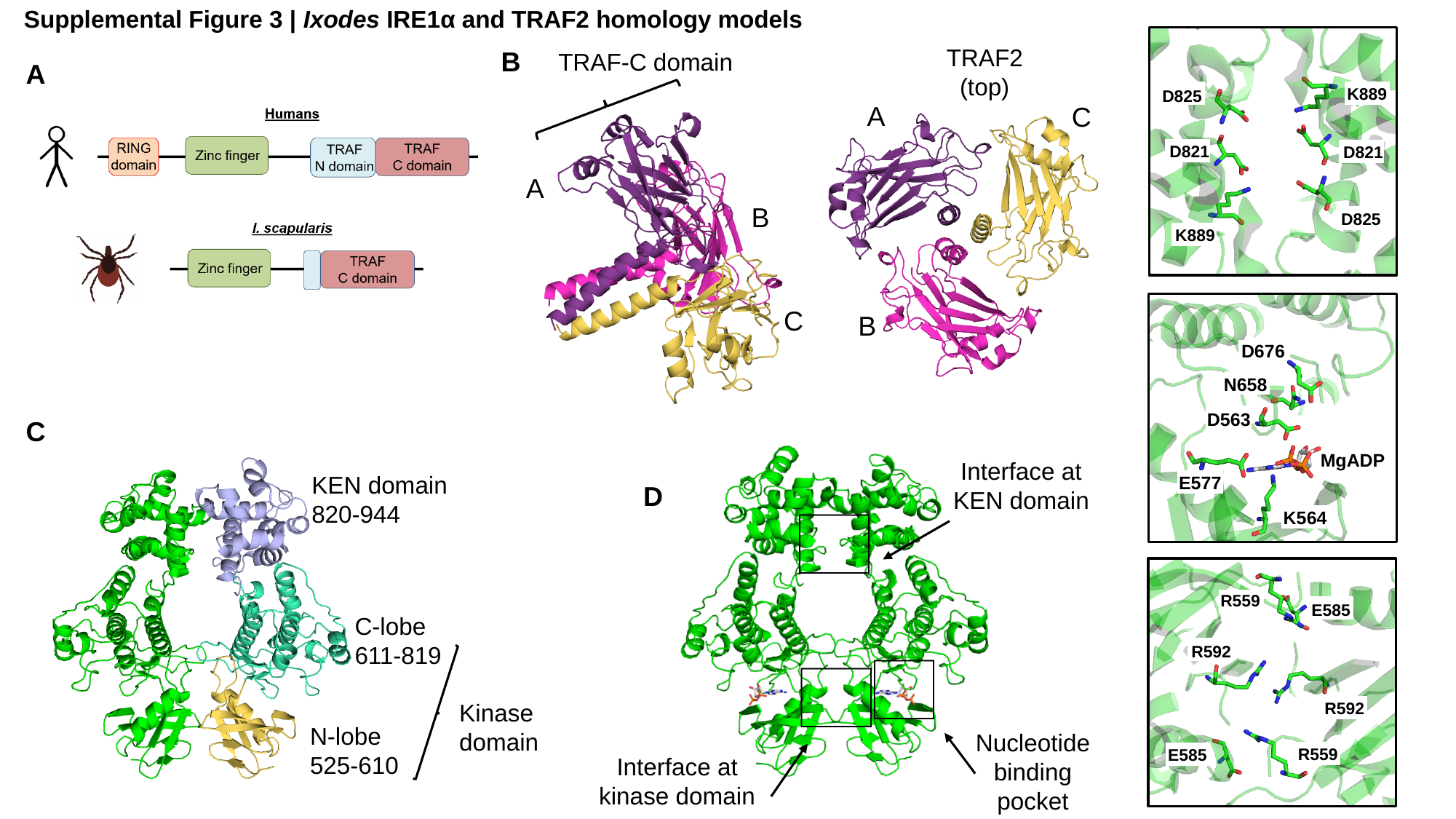

Supplemental Figure 3 | Ixodes IRE1α and TRAF2 homology models
K889
D825
D821
D821
D825
K889
B
D676
N658
D563
MgADP
E577
K564
R559
E585
R592
R592
R559
E585
TRAF2
(top)
TRAF-C domain
A
A
C
A
B
C
B
C
Interface at KEN domain
KEN domain
820-944
D
C-lobe
611-819
Kinase domain
N-lobe
525-610
Nucleotide binding pocket
Interface at kinase domain

### Slide 5
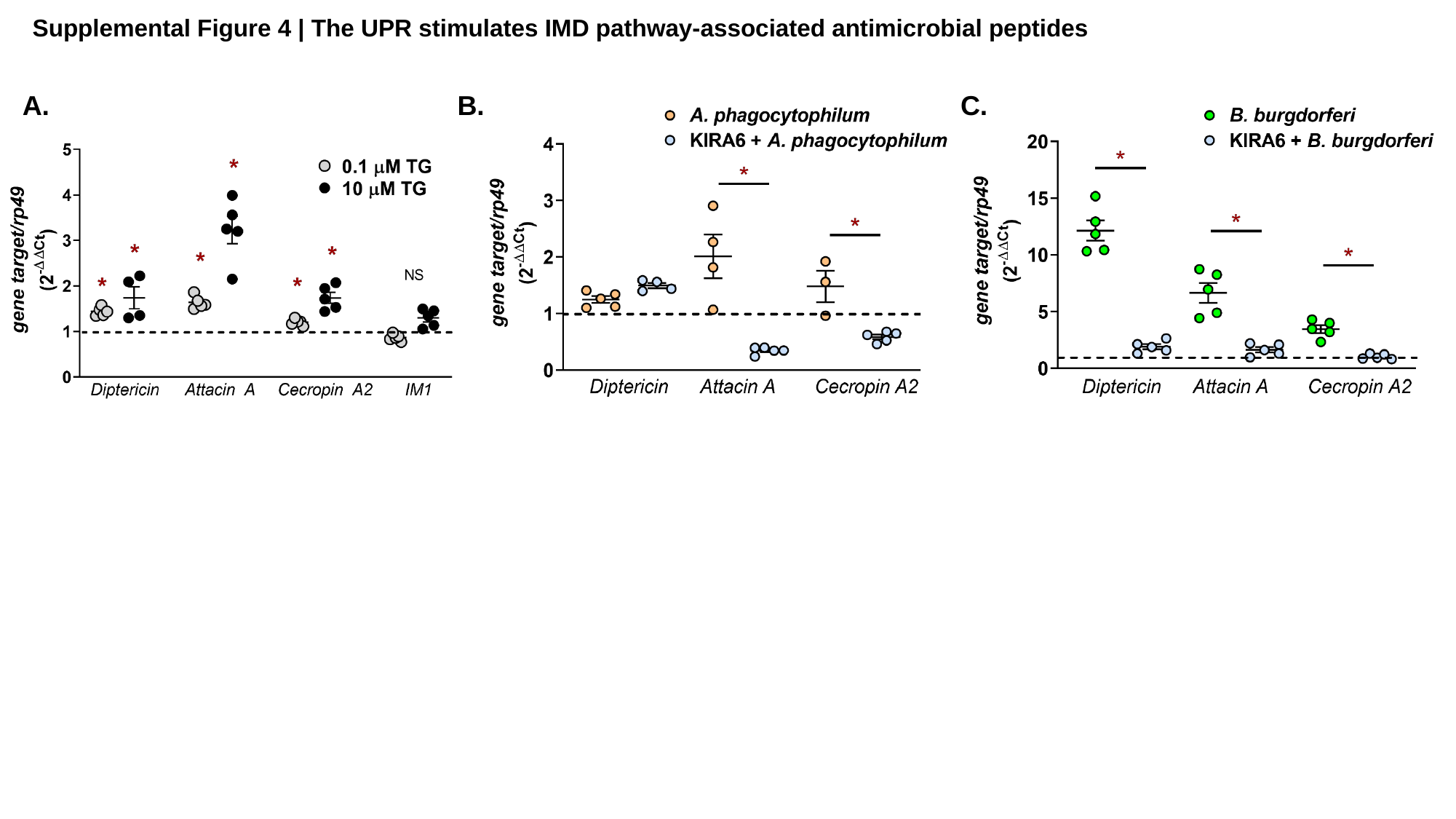

Supplemental Figure 4 | The UPR stimulates IMD pathway-associated antimicrobial peptides
A.
B.
C.

### Slide 6
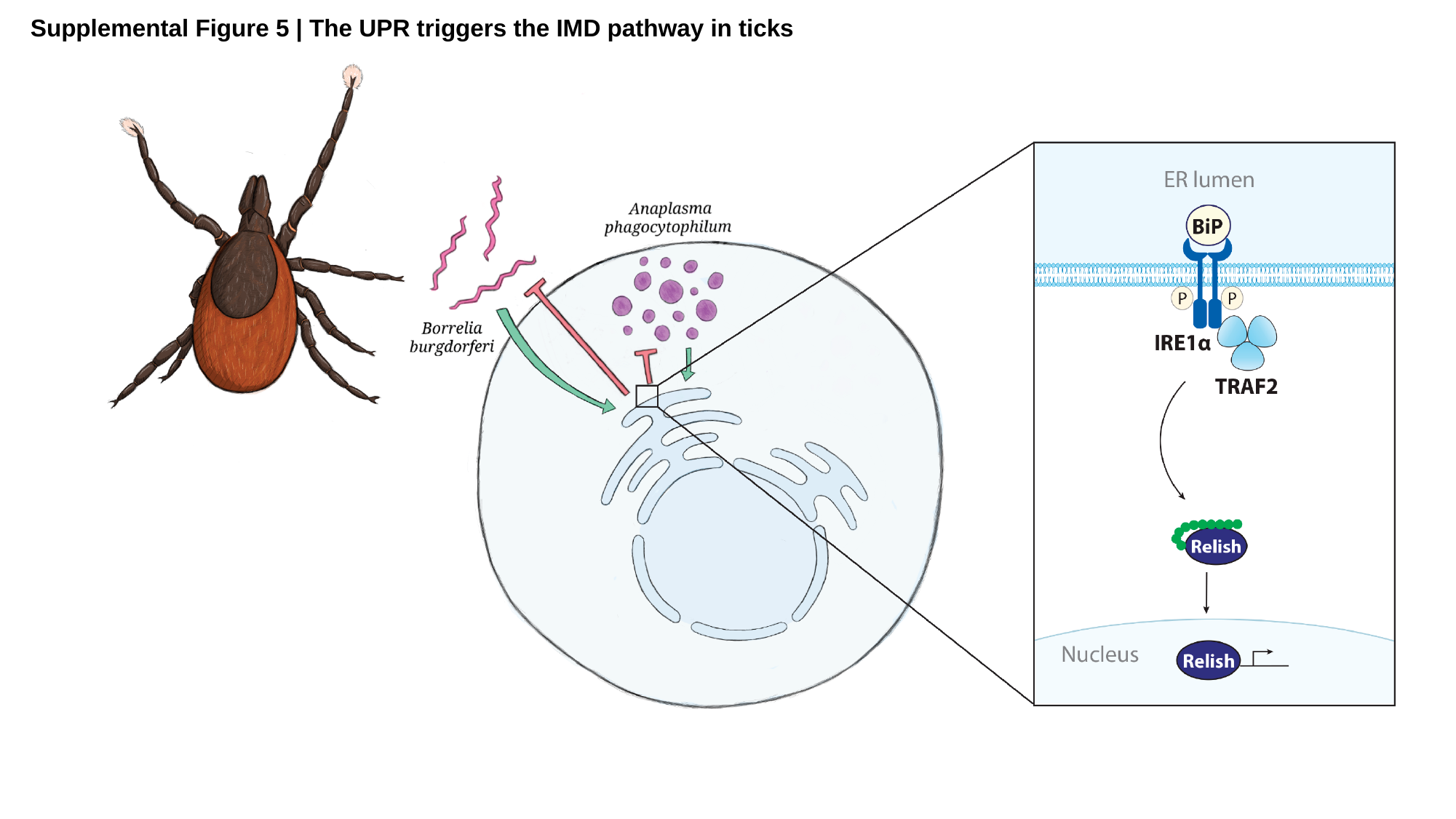

Supplemental Figure 5 | The UPR triggers the IMD pathway in ticks
